# Prime editor-based high-throughput screening reveals functional synonymous mutations in the human genome

**DOI:** 10.1101/2024.06.16.599253

**Authors:** Xuran Niu, Wei Tang, Yongshuo Liu, Ying Liu, Binrui Mo, Ying Yu, Wensheng Wei

**Author notes:** These authors contributed equally (Xuran Niu, Wei Tang, Yongshuo Liu, Ying Liu).

## Abstract

Synonymous mutations are generally considered neutral, while their roles in the human genome remain largely unexplored. Herein, we employed the PEmax system to create a library of 297,900 epegRNAs and performed extensive screening to identify synonymous mutations that impact cell fitness. While most synonymous mutations are neutral, we found that some can elicit phenotypic effects. By developing a specialized machine learning tool, we uncovered their impact on various biological processes such as mRNA splicing and transcription, supported by multifaceted experimental evidence. Notably, synonymous mutations can alter RNA folding and affect translation, as demonstrated by PLK1_S2. By integrating screening data with our model, we successfully predicted clinically deleterious synonymous mutations. This research deepens our understanding of synonymous mutations and provides insights for clinical disease studies.

## Main

Due to the degeneracy of the genetic codon system, not all single-base mutations lead to changes in the amino acid sequence, such mutations are termed synonymous and are traditionally viewed as neutral in evolutionary theory ^1^. However, a recent study in *Saccharomyces cerevisiae* challenges this traditional perception by demonstrating that both synonymous and nonsynonymous mutations can disrupt cell fitness to a similar extent, indicating a non-neutral effect ^2^. Earlier studies in viruses and prokaryotes have also suggested that synonymous mutations could affect the fitness of these organisms ^3–6^. Yet, it remains unclear whether these findings in non-eukaryotic organisms and yeast are applicable to mammals, especially humans.

Previous research has linked a small number of synonymous mutations to human diseases ^7^ and identified them as potential drivers in cancer through bioinformatics analyses ^8^. Despite the development of tools for predicting deleterious synonymous mutations ^9,10^, experimentally confirmed cases in humans remain scarce, highlighting the need for a standardized experimental method for large-scale studies of synonymous mutations in human cells.

The advent of CRISPR/Cas gene-editing technology has provided a powerful tool for studying the genome ^11,12^. An important derivative of CRISPR/Cas, the prime editor (PE), combines dCas9 with a reverse transcriptase to introduce various types of edits into the genome through a reverse transcription template in the pegRNA ^13^. Enhancements such as epegRNA ^14^ and the more efficient PEmax ^15^ have further improved the utility of PE for precise genetic manipulations. In this study, using PE technology, we created a library of epegRNAs targeting 3,644 human protein-coding genes to screen for potentially functional synonymous mutations affecting human cell fitness. Our findings confirm that while most synonymous mutations in human are neutral, a minority can produce phenotypic changes. Utilizing machine learning, we identified how these mutations influence a range of biological processes, including mRNA splicing, folding, transcription, and translation. Furthermore, we also predicted clinically deleterious synonymous mutations, thereby enhancing our understanding of these mutations and their significance in clinical research on human diseases.

### Development of a screen method for functional synonymous mutations using the prime editor

To precisely and efficiently generate synonymous mutations across different types of nucleotide substitutions, we utilized the PEmax system alongside epegRNA for targeted genetic edits. To facilitate high-throughput screening at a high multiplicity of infection (MOI), we integrated barcodes into the external region of the epegRNA, termed eBARs, similarly as we reported previously ^16,17^. Each epegRNA was labelled with three independent eBARs, effectively creating three biological replicates for our screening process (Extended Data Fig. 1a). For the screening, we selected the human colon cancer cell line HCT116, which is beneficial for PE editing due to its naturally homozygous nonsense mutation in the *MLH1* gene ^15^.

Following a set of specific criteria (Extended Data Fig. 1b), we constructed an epegRNA library. Initially, we sourced potential pathogenic synonymous mutations from two human disease databases, ClinVar and SynMICdb ^18^. Additionally, our goal was to investigate the function of synonymous mutations in a relatively unbiased manner, rather than focusing solely on clinically recognized mutations. Consequently, we selected 67 genes for saturated synonymous mutation design based on their mutation load and expression levels, including human homologous genes previously studied in yeast ^2^ (Extended Data Fig. 1c). Within this group, 11 essential human genes were selected for complete saturation tiling mutation design to thoroughly evaluate the biological impacts of synonymous mutations alongside various nonsynonymous mutations (Extended Data Fig. 1c). The library also included designs for single base insertions or the introduction of premature stop codons for gene knockouts, and incorporated *AAVS1*-targeting and nontargeting epegRNAs as negative controls. Each mutation site was targeted by an average of 2.2 epegRNAs (Extended Data Fig. 1d). For the 11 genes designed for saturation mutagenesis, all possible types of amino acid substitutions within the range of point mutations were included (Extended Data Fig. 1e). Ultimately, the library contained 297,900 epegRNAs targeting 94,993 synonymous mutations and 39,336 nonsynonymous mutations across 3,644 protein-coding genes (Extended Data Table 1 and Supplementary Table 1 and 2).

PEmax was stably integrated into HCT116 cells through lentiviral transduction, and a single clone was selected for subsequent studies. The expression profiles of HCT116-PEmax cells were almost identical to those of wild-type cells and the HCT116-PEmax cells infected with nontargeting or *AAVS1*-targeting epegRNAs, which served as negative controls, showed minimal changes (Extended Data Fig. 2a). This indicates that the stable cell line closely resembles the characteristics of wild-type HCT116 cells. We also assessed the editing efficiency of PEmax in HCT116 cells. Over a 28-day culture period with periodic sampling, the editing of the high-efficiency *FANCF* +5 G to T mutation showed a gradual slowdown after 14 days, whereas the low-efficiency *RNF2* +1 C to A mutation exhibited a gradual increase in editing efficiency over time (Extended Data Fig. 2b). Given that phenotypic changes often lag behind genotypic alterations, we extended the screening duration to 35 days. We termed this methodology PRESENT (<u>PR</u>ime <u>E</u>ditor based <u>S</u>cre<u>EN</u> <u>T</u>echnology) (Fig. 1a).

**Fig. 1.**
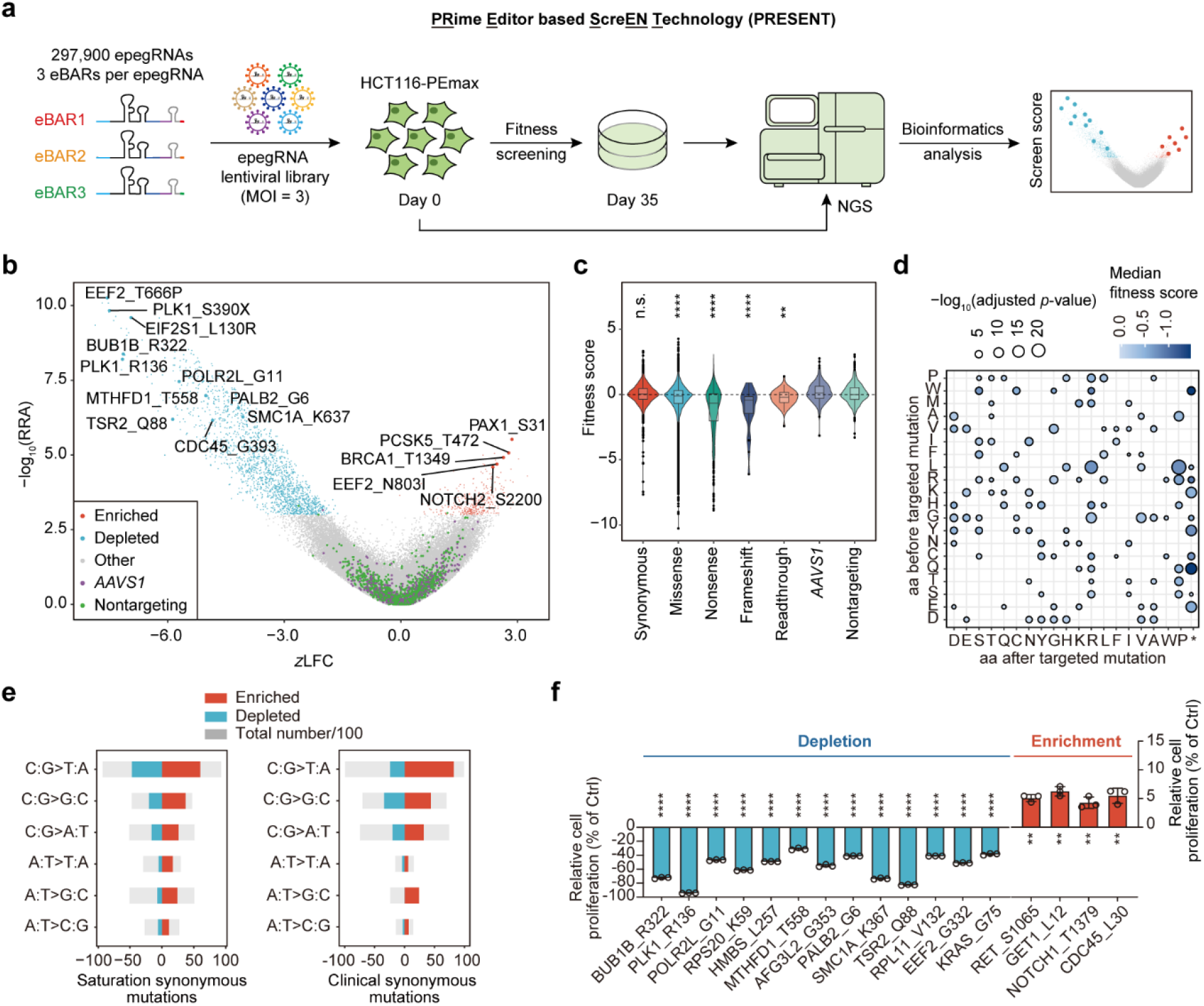
High-throughput screening unveils functional synonymous mutations in the human genome. **a**, Schematic of the PRESENT workflow. **b**, Volcano plot illustrating the results of screening for functional synonymous and nonsynonymous mutations affecting cell fitness. Blue and red dots denote depleted and enriched epegRNAs, respectively. **c**, Distribution of synonymous and various nonsynonymous mutations in the cell fitness screen. *P* value was calculated following Benjamini-Hochberg correction. \*\**P* < 0.01, \*\*\*\**P* < 0.0001; n.s., not significant. **d**, Effects of different amino acid (aa) substitutions on cell fitness. **e**, Distribution of mutations enriched across different types of CG and AT base pair substitutions in both saturation synonymous mutations and clinical synonymous mutations. **f**, Relative cell proliferation rates for validated mutations. The data are presented as mean ± SD (n = 3). *P* values were calculated using Student’s *t* test, \*\**P* < 0.01, \*\*\*\**P* < 0.0001; n.s., not significant.

### High-throughput screening reveals the impact of mutations on cell fitness

Since mutations are introduced through the reverse transcription template (RTT), decoding the information from the RTT and eBAR regions is sufficient (Extended Data Fig. 3a). The internal replicates represented by the three eBARs were well correlated throughout the entire screening process (Extended Data Fig. 3b, c). We devised a bioinformatics algorithm tailored specifically for this screen (named ZFC-eBAR, see Methods) and established a screen score threshold of 3 based on the performance of the negative controls. A total of 1,917 mutations impacting cell fitness were identified, showing trends of both depletion and enrichment (Fig. 1b and Supplementary Table 3). Among these, 409 were synonymous mutations, and 1,508 were nonsynonymous mutations. In our screen, the occurrence rate of synonymous mutations was only 0.38%, compared to 5.32% for nonsynonymous mutations. While the fitness impact of synonymous mutations didn’t significantly differ from that of the negative controls, nonsynonymous mutations, including missense and nonsense types, exhibited clear effects on cell fitness (Fig. 1c). This indicates that, in general, synonymous mutations in the human genome are neutral, diverging from results seen in yeast ^2^. Significant fitness differences between synonymous and missense mutations were noted within the 11 genes targeted for saturated mutation design (Extended Data Fig. 4a). Further analysis of these nonsynonymous mutations demonstrated a direct relationship between the probability of amino acid substitution and the effect on cell fitness. Amino acids with lower substitution probabilities are more likely to exhibit more significant fitness impacts after mutation (Extended Data Fig. 4b). Nonsense mutations typically produced the most pronounced biological effects, with amino acid substitutions such as L>P, L>R, R>P more frequently resulting in phenotypes (Fig. 1d and Extended Data Fig. 4c).

Although synonymous mutations in the human genome are predominantly neutral, the 409 enriched mutations identified in our study demonstrated biological effects. Statistical analysis of these mutations revealed that the highest levels of enrichment occurred in alanine, glycine, and leucine following synonymous changes. Nevertheless, the observed patterns differed when compared to those in clinical synonymous mutations (Extended Data Fig. 4d). Furthermore, CG base pairs in these synonymous mutations were more likely to generate fitness effects post-mutation compared to AT base pairs, a trend consistent with both saturation synonymous mutations and clinical synonymous mutations (Fig. 1e). This consistency exists regardless of the preferences associated with prime editing^19,20^.

We further validated the impact of these enriched synonymous mutations on cell proliferation (Fig. 1f and Extended Data Fig. 5a-d). All tested mutations showed precise and clean editing results (Extended Data Fig. 5e). Over an 18-day period of continuous cell culture, significant phenotypic effects were observed for all synonymous mutations categorized under depletion, while those under enrichment showed relatively mild effects. Based on these observations, our subsequent research efforts concentrated on exploring the effects of depletion-direction synonymous mutations.

### Machine learning analysis identifies key factors influencing deleterious synonymous mutations

To elucidate the mechanisms behind deleterious synonymous mutations, we analyzed them from three perspectives: gene, mRNA, and nucleotide level. Deleterious mutations are more likely to cause aberrant splicing, disrupt RNA secondary structure, and utilize infrequent codons. These mutations often occur in conserved nucleotides, highly expressed genes, and essential genes (Extended Data Fig. 6a-f).

In order to identify the most influential features associated with these mutations, we developed a machine learning model named DS Finder (<u>D</u>eleterious <u>S</u>ynonymous mutations Finder) based on the CatBoost framework ^21^ and trained it as a binary classifier with our screening data (Fig. 2a). Given the extensive data from our screen, DS Finder outperformed two existing models: CADD, which predicts general mutations ^10^, and SilVA, a predictor for synonymous mutations trained with a smaller dataset of 33 deleterious synonymous mutations ^9^ (Fig. 2b).

**Fig. 2.**
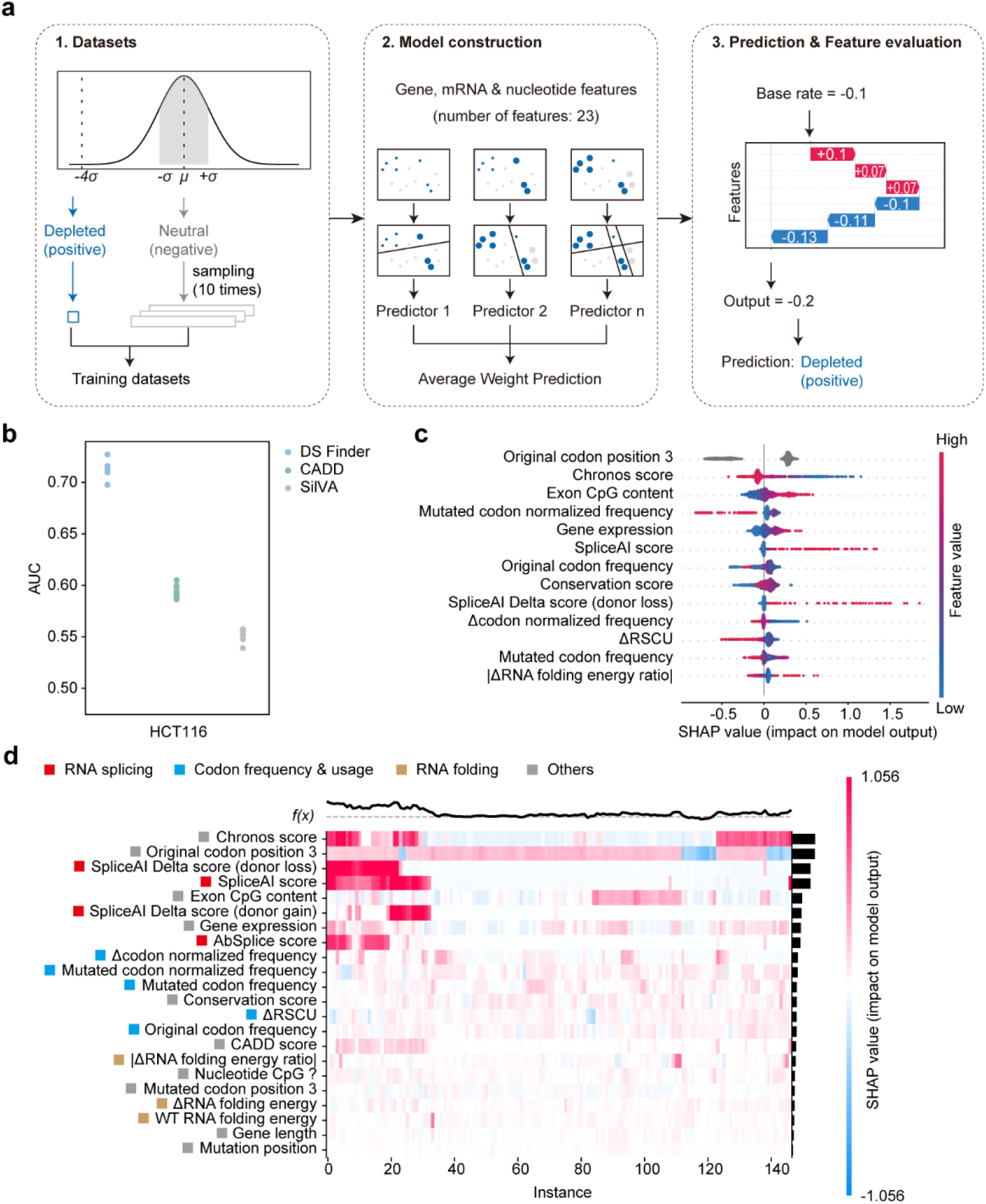
Machine learning model for analyzing determinants of deleterious synonymous mutations. **a**, Workflow diagram of DS Finder. The dataset was selected using a 1:10 ratio. The model was developed using CatBoost, and SHAP (SHapley Additive exPlanations) was used to determine feature importance. **b**, Performance of DS Finder in the HCT116 cell line compared with CADD and SilVA. Each point on the graph represents a different dataset, totaling 10. **c**, The relative importance of various features in predicting the effects of synonymous mutations. **d**, Heatmap of SHAP values, illustrating the supervised clustering method that categorizes data points based on their feature explanations.

Key features identified by DS Finder include the third position of the codon being CG or AT base pairs, with CG pairs are more likely to impact cell fitness (Fig. 1e and Fig. 2c). Factors like gene essentiality, expression levels, and exon CpG content also play crucial roles (Fig. 2c), emphasizing the need to consider these features in the model as they vary across different tissues. Alterations in splicing, especially the loss of splice donor sites (Fig. 2c), likely representing the primary mechanism driving deleterious effects (Fig. 2d). Moreover, synonymous mutations can adversely affect cell fitness by altering codon usage and mRNA folding (Fig. 2c, d).

Altogether, our machine learning model shows that the deleterious effects of synonymous mutations are driven by a combination of factors including splicing disruptions, codon bias, and nucleotide conservation.

### Synonymous mutations generated aberrant splicing events

DS Finder identified disruption of splicing as a key mechanism through which synonymous mutations exert their deleterious effects. SpliceAI ^22^, a widely used tool for predicting erroneous splicing events, revealed that 23.13% of synonymous mutations in the depletion direction could be deleterious, affecting essential gene expression (Extended Data Fig. 7a). Most of these mutations alter splice donor sites, with a few mutations, such as KIF11_A133, introducing new splice acceptor in transcripts (Extended Data Fig. 7b-e). Mutations causing splice donor loss accounted for 12% of the total, followed by mutations that create new splice donors, which accounted for 7% (Fig. 3a, b).

**Fig. 3.**
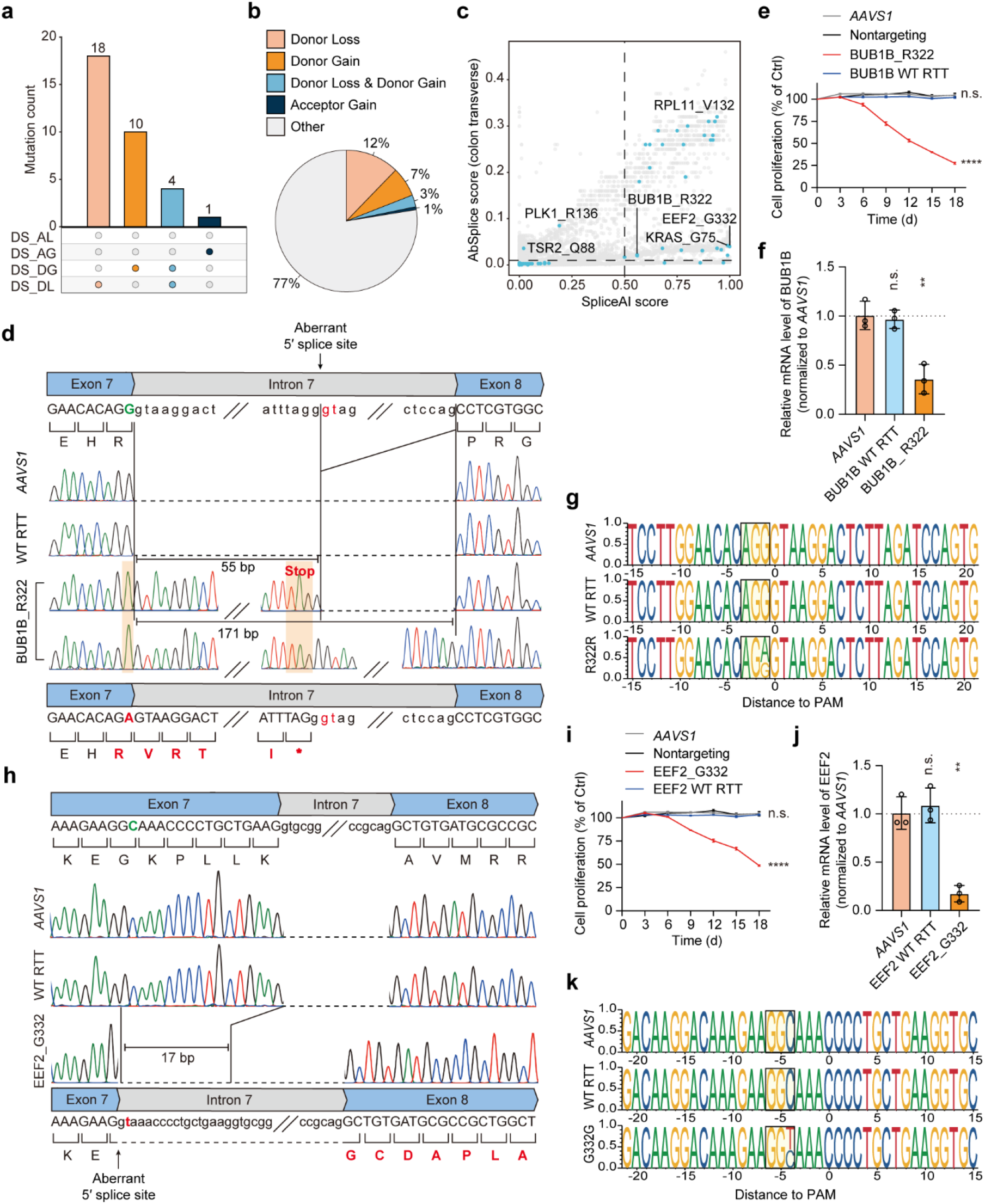
Impact of synonymous mutations on aberrant RNA splicing. **a-b**, Statistical analysis of synonymous mutations leading to various aberrant RNA splicing events. **c**, Integration of predictions from SpliceAI and AbSplice tools. **d**, Schematic depiction of the splicing alterations caused by the BUB1B_R322 (AGG>AGA) mutation. **e**, Validation of the effect of the BUB1B_R322 (AGG>AGA) mutation on cell proliferation. **f**, Relative mRNA expression levels of *BUB1B* in experimental and control groups. The mRNA level of each sample was quantified by real-time qPCR and normalized by *GAPDH*, and the indicated relative mRNA level was normalized to that of *AAVS1*-targeting control cells. **g**, Analysis of editing outcomes for epegRNA targeting BUB1B_R322 and controls via NGS. **h**, Schematic depiction of the splicing alterations caused by the EEF2_G332 (GGC>GGT) mutation. **i**, Validation of the effect of the EEF2_G332 (GGC>GGT) mutation on cell proliferation. **j**, Relative mRNA expression levels of *EEF2* in the experimental and control groups. The mRNA level of each sample was quantified by real-time qPCR and normalized by *GAPDH*, and the indicated relative mRNA level was normalized to that of *AAVS1*-targeting control cells. **k**, Analysis of editing outcomes for epegRNA targeting EEF2_G332 and controls via NGS. All data are presented as mean ± SD (n = 3). *P* values were calculated using Student’s *t* test, \*\**P* < 0.01, \*\*\*\**P* < 0.0001; n.s., not significant.

Additionally, a new model called AbSplice has been developed to predict aberrant splicing caused by mutations in specific human tissues ^23^. Using AbSplice, we predicted new mutations, such as PLK1_R136 (AGG>AGA) and TSR2_Q88 (CAG>CAA), that could lead to aberrant splicing in colon tissue (Fig. 3c). Overall, nearly a quarter of the synonymous mutations from the depletion screening could potentially lead to aberrant splicing events.

We selected a subset of these mutations for validation, using epegRNAs with reverse transcription templates matching the reference sequence at each mutation site as controls to eliminate editing process effects. The BUB1B_R322 (AGG>AGA) mutation, a top-ranked synonymous mutation from the depletion results (Fig. 1b) located at the junction of exon 7 and intron 7 of *BUB1B*, disrupts the original splice donor site (Donor Loss type), resulting in intron retention (Fig. 3d). Interestingly, this mutation led to two abnormal transcript variants: one with complete intron 7 retention and another with a newly created splice donor site within intron 7, leading to a truncated intron retention (Fig. 3d). Both variants could lead to mRNA degradation due to the generated premature stop codon. We further validated the impact of this mutation on cell fitness and mRNA abundance (Fig. 3e-g). Furthermore, BUB1B_R322 (AGG>AGA) is also noted in ClinVar in patients with mosaic variegated aneuploidy syndrome 1 but labeled with “uncertain significance,” highlighting our screen can reveal synonymous mutations that may be misannotated in clinical databases. Another mutation, RPL11_V132 (GTG>GTC), disrupts a splice donor, causing retention of the intronic region and altering cellular phenotypes (Extended Data Fig. 8a-d).

EEF2_G332 (GGC>GGT) represents a different impact on splicing by generating a new splice donor within an exon (Donor Gain type). This new splice donor precedes the original, resulting in partial exon excision and a frameshift in the downstream sequence (Fig. 3h). This abnormal transcript significantly reduces cell fitness and mRNA levels (Fig. 3i-k). A similar case is observed in KRAS_G75 (GGG>GGT) (Extended Data Fig. 8e-h). These findings suggest synonymous mutations near exon-intron boundaries can induce aberrant splicing, affecting gene expression and cellular phenotypes.

### Synonymous mutations impacted RNA stability and influenced translation processes

In addition to aberrant RNA splicing, synonymous mutations can also impact RNA folding and translation within cells (Fig. 2d). A specific mutation, PLK1_S2 (AGT>AGC), found early in the coding sequence of the essential gene *PLK1*, enhanced RNA stability (Fig. 4a). Despite both codons encoding serine, AGC is used more frequently than AGT (codon frequencies of 19.5 per thousand vs. 12.1 per thousand, respectively), implying that changes in cell fitness might stem from alterations in mRNA folding rather than codon usage. Predicted analysis of mRNA structure pre- and post-mutation revealed that the PLK1_S2 (AGT>AGC) mutation enhanced the stability of the local mRNA structure near the start codon (Fig. 4b and Extended Data Fig. 9a-c).

**Fig. 4.**
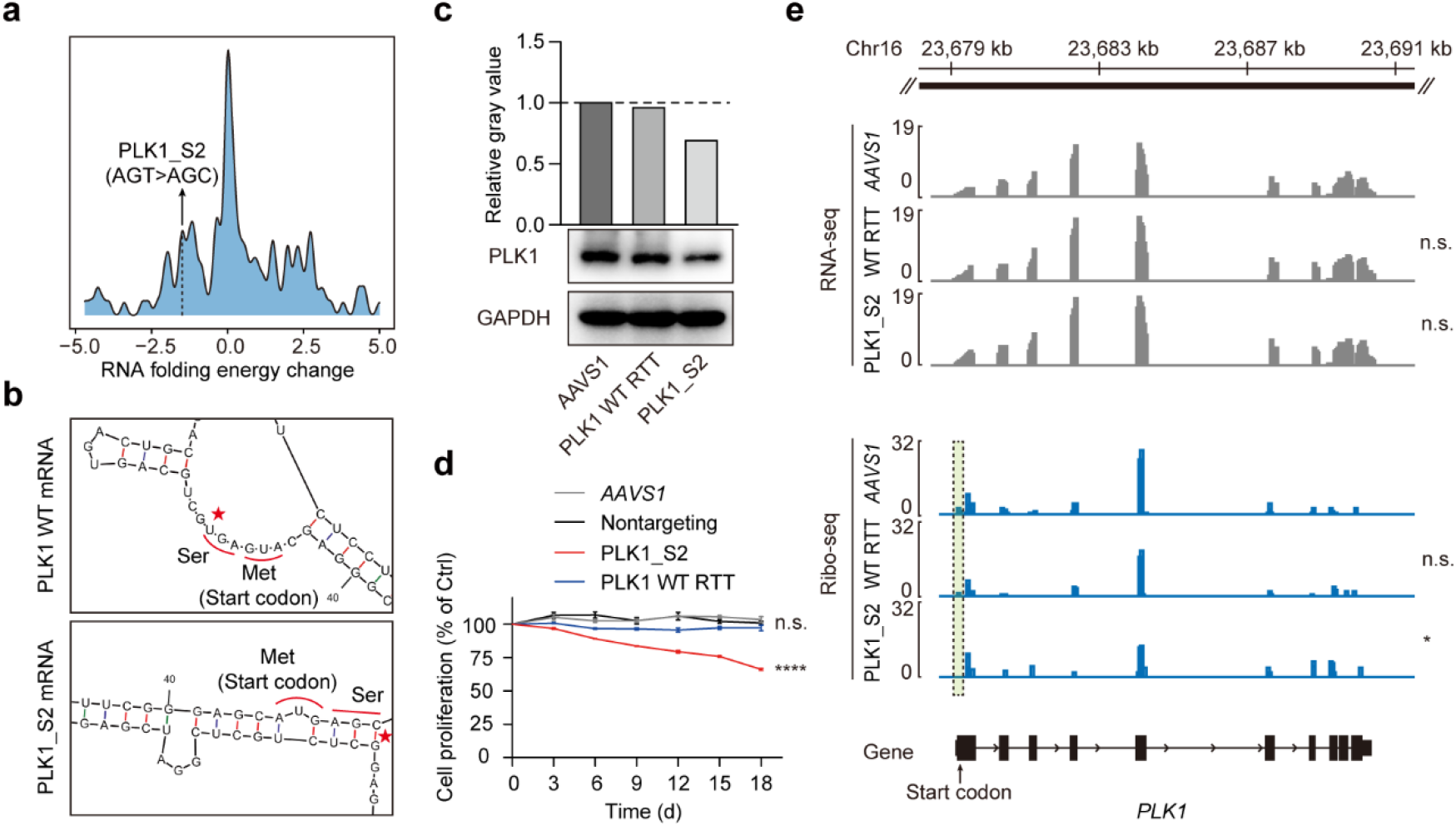
Influence of synonymous mutations on protein translation through RNA folding. **a**, Mapping of RNA folding energy changes for synonymous mutations in the depletion direction. **b**, Comparison of local mRNA structure of *PLK1* before and after the PLK1_S2 (AGT>AGC) mutation, predicted using RNA Folding Form. **c**, Western blot analysis comparing PLK1 protein levels between PLK1_S2 and controls. **d**, Validation of the effect of the PLK1_S2 (AGT>AGC) mutation on cell proliferation. Data are presented as mean ± SD (n = 3). *P* values were calculated using Student’s *t* test, \*\*\*\**P* < 0.0001; n.s., not significant. **e**, RNA-seq and Ribo-seq analyses of PLK1_S2 and control, with two biological replicates per group. *P* values were calculated using Student’s *t* test, \**P* < 0.05; n.s., not significant.

To verify if this mutation affected *PLK1* expression, we conducted a western blot analysis to measure PLK1 protein levels. Results indicated that the wild-type reverse transcription template (WT RTT) had no impact on PLK1 protein levels, whereas the PLK1_S2 (AGT>AGC) mutation led to decreased protein abundance (Fig. 4c). Additionally, this mutation did not impact on neighboring codons either upstream or downstream of the mutation site (Extended Data Fig. 10a), and the observed decrease in cell fitness was solely attributable to this single synonymous mutation (Fig. 4d).

Considering that the mutation-induced stem structure disrupted the original loose structure near the start codon of WT PLK1 mRNA (Fig. 4b), potentially impeding translation initiation and conferring a translation disadvantage, we further used Ribo-seq to assess translation at this site. The results indicated that PLK1_S2 (AGT>AGC) mutation made ribosome binding at the start codon more challenging, without affecting transcription levels (Fig. 4e and Extended Data Fig. 10b). The difficulty in initiating translation consequently led to reduced protein expression. This scenario underscores the relationship between changes in RNA folding induced by synonymous mutations and their effects on the translation process, ultimately influencing cellular functions through altered protein levels.

### Single-cell sequencing used to investigate the effects of synonymous mutations on gene expression

Through these studies, we have elucidated potential mechanisms through which functional synonymous mutations influence biological processes. Recognizing prior studies that suggest synonymous mutations can alter intracellular RNA abundance ^2^, we aimed to systematically assess the impact of synonymous mutations identified from our screening on gene expression at a high-throughput level. To achieve this, we integrated a single-cell screening approach with our PRESENT, calling it DIRECTED-seq (<u>DIR</u>ect <u>E</u>pegRNA <u>C</u>apture and <u>T</u>arget<u>ED</u> <u>seq</u>uencing). We constructed a DIRECTED-seq epegRNA library targeting each synonymous mutation using the prime editor to systematically investigate their effects on gene expression. We selected nearly all synonymous mutations that were either enriched or depleted, excluding those within genes expressed at low levels, resulting in a total of 370 mutations. Additionally, the library also included several negative controls: 15 nontargeting epegRNAs, 15 *AAVS1*-targeting epegRNAs, and 10 epegRNAs targeting synonymous mutations that were not enriched.

HCT116-PEmax cells were transduced with this pooled library at a high MOI of about 5, followed by selection with puromycin. After 14 days of culture, we simultaneously captured the epegRNAs and transcriptomes from single cells (Fig. 5a). A custom primer designed from the evopreQ1 motif achieved the reverse transcription of epegRNAs, and the transcriptome library was generated using an oligo-dT reverse transcription primer (Extended Data Fig. 11a). Furthermore, bulk RNA-seq was performed on cells from the same batch to evaluate the editing efficiency (Fig. 5a). The majority of epegRNAs were effective, showing an average editing efficiency of about 15% for mutations detectable at sufficient sequencing depth (Fig. 5b).

**Fig. 5.**
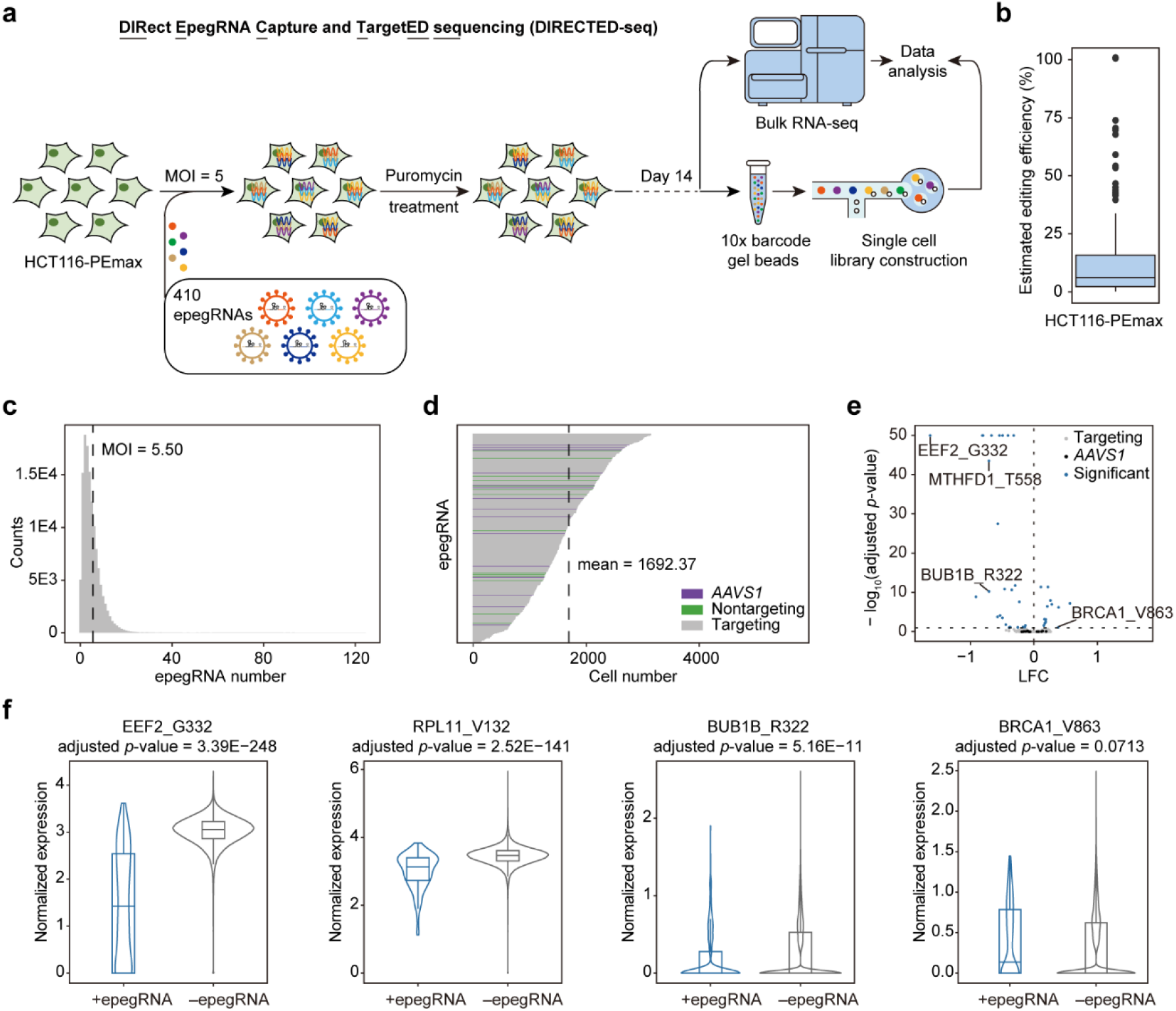
Investigating the impact of synonymous mutations on gene expression using DIRECTED-seq. **a**, Workflow diagram for DIRECTED-seq, conducted in HCT116-PEmax cells. Cells were transduced with the epegRNA library at a MOI of 5 and subsequently treated with puromycin. On Day 14, both bulk RNA sequencing and single-cell sequencing were performed. **b**, Estimated editing efficiency at sites where mutations were expected to be detectable. **c**, Histogram showing the distribution of the number of epegRNAs per cell, with the mean value indicated by a dashed line. **d**, Distribution of the number of cells detected for each epegRNA, categorized by coverage. The dashed line represents the mean value, with purple, green, and gray lines representing *AAVS1*-targeting, nontargeting, and mutation-targeting epegRNAs, respectively. **e**, Volcano plot displaying the effects of synonymous mutations on the expression of corresponding genes. **f**, Single-cell gene expression profiles for cells harboring epegRNAs targeting EEF2_G332, RPL11_V132, BUB1B_R322, and BRCA1_V863. *P* values were calculated using the SCEPTR method and adjusted with the Benjamini-Hochberg correction.

We recovered 129,193 single cells, averaging 5.5 epegRNAs per cell, with each targeted synonymous mutation represented in approximately 1,692 cells (Fig. 5c, d). Consistent with previous findings ^24^, a positive correlation was observed between the number of epegRNAs and UMIs per cell (Extended Data Fig. 11b), suggesting sequencing depth as a potential confounding factor. To address this, we employed a conditional resampling approach (SCEPTRE) for our differential expression analysis ^24^. Nontargeting epegRNAs served as negative controls, showing no observable effects (Extended Data Fig. 11c, d). Ultimately, we identified 40 synonymous mutations that significantly influenced gene expression, at a 10% false discovery rate (FDR) (Fig. 5e and Supplementary Table 7).

A notable finding was the substantial decrease in *EEF2* gene expression caused by the synonymous mutation EEF2_G332 (GGC>GGT), where expression dropped to 32% of its original level (log2FC= -1.63, adjusted *p*-value=3.39E-248). Given EEF2’s essential role in cell fitness, this significant reduction correlates with the mutation’s presence in the depletion direction of the screen. Similar effects were observed with synonymous mutations in other essential genes. For instance, substitution of GTG with GTC, GTT, and GTA at RPL11_V132 (log2FC = -0.488, -0.312, and -0.401, respectively, with adjusted *p*-values of 2.52E-141, 2.36E-135, and 8.33E-77, respectively) and substitution of AGG with AGA at BUB1B_R322 (log2FC= -0.702, adjusted *p*-value=5.16E-11) significantly lowered gene expression, which corresponded with documented aberrant splicing events (Fig. 3d-k and Extended Data Fig. 8a-d). Conversely, some synonymous mutations, such as BRCA1_V863 (GTT>GTG), were found to increase gene expression (log2FC = 0.389, adjusted *p*-value = 0.0713). Given BRCA1’s critical roles in DNA damage repair, cell cycle checkpoint control, and genomic stability maintenance, these mutations might substantially impact cellular functions and tumor suppression mechanisms (Fig. 5f and Supplementary Table 7).

These results from DIRECTED-seq effectively link specific mutations to changes in gene expression, unveiling their potential biological impacts. Although changes in transcript abundance could be driven by several mechanisms, such as misregulated RNA splicing, not all are linked to such changes. For instance, BRCA1_V863 (GTT>GTG) may affect RNA stability independently of splicing disruptions (Supplementary Table 6). Mutations could destabilize the global RNA structure, leading to increased transcript degradation, or conversely, they may enhance the stability of transcripts^25^. Additionally, transcripts containing infrequently used codons during the early stage of translation elongation, referred to as the ramp, may also be subjected to degradation due to the slow translation speed of such transcripts ^26,27^. These potential impacts highlight the need for further studies to explore how mutations influence transcript abundance and their broader biological effects.

### Identifying new deleterious synonymous mutations linked to disease risk

We have systematically studied functional synonymous mutations in our library using PRESENT and DIRECTED-seq. However, relying solely on screening and experimentation to uncover potential deleterious synonymous mutations in clinical databases has its limitations. To overcome this, we employed our novel machine learning model, DS Finder, trained on our screening data, to identify novel functional clinical mutations across a broader dataset (Fig. 6a). We focused on mutations related to colonic diseases in the ClinVar database, such as lactose intolerance and inflammatory bowel disease, which share similar genetic backgrounds to the HCT116 cell line used in our studies. Through DS Finder predictions, we identified several potentially deleterious synonymous mutations (Supplementary Table 9).

**Fig. 6.**
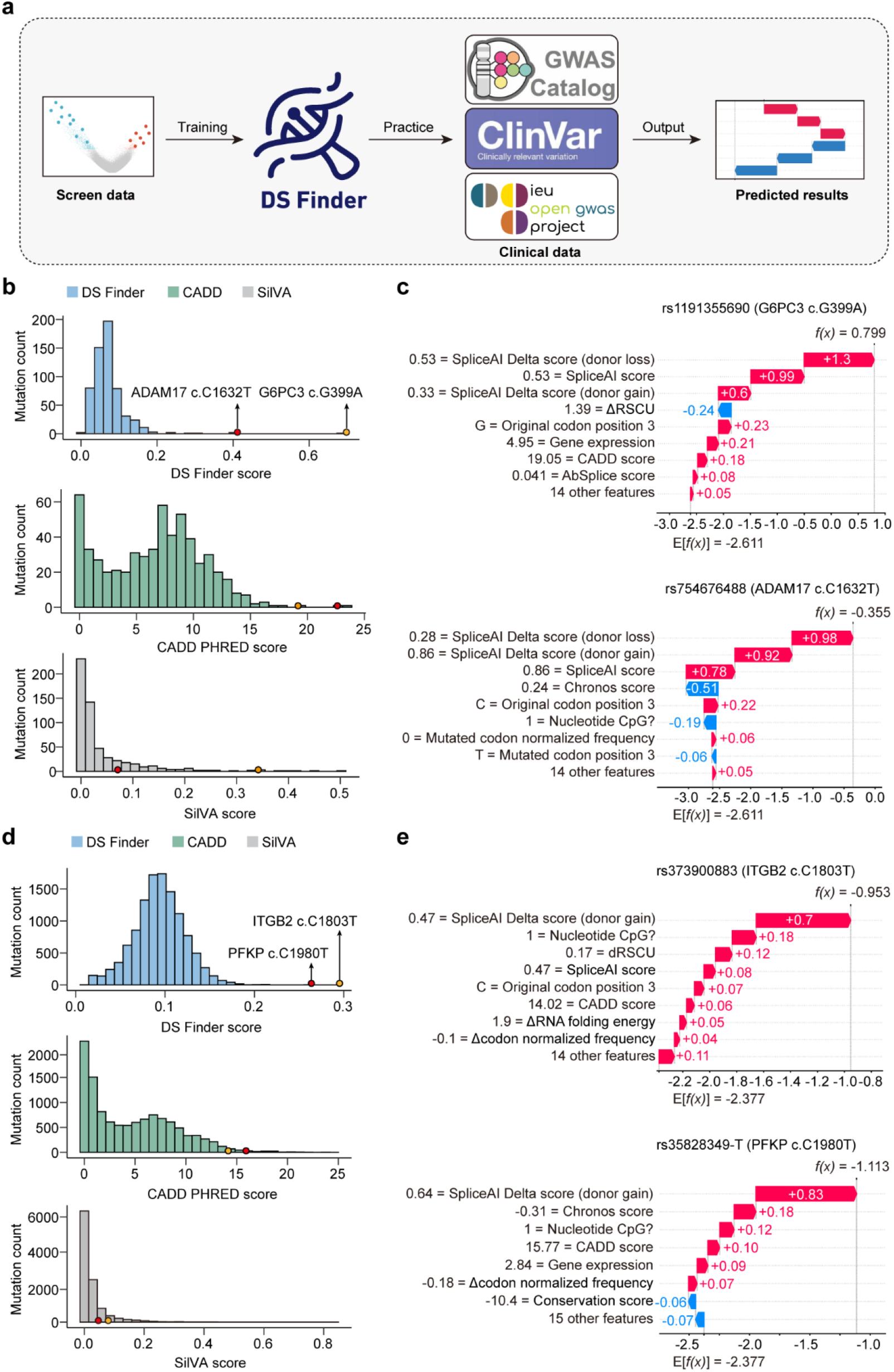
Predicting deleterious synonymous mutations in clinical databases. **a**, Flowchart illustrating the process of training the DS Finder using screening data and its application to clinical data for identifying novel deleterious synonymous mutations. **b**, Prediction scores from three models for synonymous mutations associated with colonic diseases. **c**, SHAP value waterfall plots for two colonic disease examples: G6PC3 c.G399A and ADAM17 c.C1632T, highlighting how individual features influence the prediction outcomes. **d**, Prediction scores from three models for synonymous mutations associated with blood diseases and relevant GWAS phenotypes. **e**, SHAP value waterfall plots for two examples related to blood diseases and relevant GWAS phenotypes: ITGB2 c.C1803T and PFKP c.C1980T, highlighting how individual features influence the prediction outcomes. In these waterfall plots, red indicates features that contribute positively to the prediction outcome, while blue indicates features that negatively impact the predictions.

One such example is the G6PC3 c.G399A, associated with autosomal recessive severe congenital neutropenia, is annotated as “likely benign” in ClinVar but was predicted as highly pathogenic in our analysis and confirmed by both SilVA and CADD (Fig. 6b). This mutation impacts splicing via donor loss, affecting the gene’s normal function and aligning with the disease’s mechanism (Fig. 6c). Similarly, the ADAM17 c.C1632T mutation found in patients with inflammatory skin and bowel diseases, also annotated as “likely benign” in ClinVar, was predicted to be pathogenic by both DS Finder and CADD (Fig. 6b), likely affecting splice donor sites and impacting gene function (Fig. 6c).

To broaden our analysis beyond colonic diseases, we extended our research to include blood disorders. Considering the importance of cell genetic backgrounds, we implemented the PRESENT in the K562 cell line using the same epegRNA^eBAR^ library (Extended Data Fig. 12a and Supplementary Table 8) and developed a corresponding prediction model that outperformed both CADD and SilVA (Extended Data Fig. 12b). We searched ClinVar for mutations associated with six blood diseases and considered three GWAS phenotypes: leukocyte disorder, leukocyte count, and white blood cell count (Supplementary Table 9). The ITGB2 c.C1803T mutation, linked to leukocyte adhesion deficiency, was identified as the top-ranked mutation (Fig. 6d). The second-ranked mutation, PFKP c.C1980T, common in the population (frequency > 0.1%) and modestly correlated with white blood cell count (effect size = 0.966189, *p* = 0.036), was also predicted to disrupt splicing via splice donor gain (Fig. 6e).

Additionally, we launched a website called *Hearing Silence* (https://search-synonymous-mutations.streamlit.app/), enabling open access to screen results from two cell lines and offering the DS Finder algorithm for free use. Overall, our model effectively identified novel pathogenic synonymous mutations in clinical databases, showing potential for broad generalizability across diverse genetic disease backgrounds. Our findings also suggest that current clinical databases may underreport deleterious synonymous mutations, underscoring the value of predictive models for more precise annotations. This research underscores the potential of exploring additional functional synonymous mutations within the human genome, which is vital for understanding disease etiology and delving deeper into their molecular genetic mechanisms.

## Discussion

In this study, we developed a high-throughput screening technique called PRESENT, utilizing the prime editor to investigate functional synonymous mutations within the human genome. Overall, our findings indicate that the majority of synonymous mutations in the human genome are neutral, including those occurring in essential genes. This contrasts with previous studies in yeast, where synonymous mutations appeared more consequential. Our library, which includes human homologs of the genes studied in yeast, didn’t show significant enrichment of these mutations, potentially due to differences between haploid yeast and diploid human cells, and the greater complexity and larger intron regions in the human genome ^28^. This suggests that the effects of synonymous mutations can vary significantly with the genetic background, challenging the simplistic classification of these mutations as universally neutral or deleterious.

Among the synonymous mutations that do exhibit biological effects, changes involving CG base pairs were particularly impactful on cell fitness. This highlights the significance of codons ending in C or G (also named as GC3) in genomic structure and their specific responses to synonymous mutations ^29^. These mutations may affect CpG content within gene sequences, a feature incorporated into our machine learning model, DS Finder. Additionally, high-frequency codons often are GC3, which may relate to the stability of anticodon pairing during translation. The pronounced biological effects of GC3 mutations suggest that they are subject to negative selection, maintaining genomic stability over evolution time.

Our observations also show that synonymous mutations can alter gene expression levels, echoing findings from yeast studies ^2^. These effects are systematically analyzed in human cells using DIRECTED-seq, primarily due to aberrant splicing events on mRNA. Dominant effects were observed at splice donor sites, likely due to the more conserved sequences upstream of these sites compared to downstream of splice acceptor sites ^30^. When gene expression is not altered, translation-coupled biological mechanisms may play a role. For example, we observed that increased mRNA stability near the start codon could impede translation initiation, reducing protein expression levels, consistent with phenomena observed in prokaryotes ^31^ and various genomic studies ^32^. Changes in protein levels might also be linked to variations in codon usage frequency. In this study, we have outlined a summary of possible explanations. While there are many other potential biological mechanisms through which synonymous mutations could affect human cell functions, further experimental investigations are necessary.

Additionally, we developed DS Finder, a prediction model for deleterious synonymous mutations that outperforms existing models like SilVA and CADD. DS Finder’s training set is more extensive than that of SilVA ^9^, which includes only 33 experimentally verified deleterious mutations, and offers greater specificity than CADD ^10^, which predicts all mutation types. DS Finder also considers cell type, tissue type, and gene background in its predictions, enhancing its accuracy and utility in clinical data analysis, making it invaluable for future clinical research and diagnostics.

Overall, this study provides new insights and experimental evidence on the impact of synonymous mutations in the human genome and their potential biological mechanisms. It underscores the precision of the prime editor over traditional CRISPR/Cas and base editors for high-throughput genomic studies ^33,34^. Looking ahead, PRESENT and DIRECTED-seq provide useful tools in characterizing and exploring mechanisms behind clinical drug-resistant mutations and other genetic phenomena.

## Acknowledgments

We thank the National Center for Protein Sciences (Beijing) at Peking University for their assistance with fluorescence-activated cell sorting and analysis, especially Dr. Hongxia Lv and Ms. Huan Yang for their technical support. We thank the High-performance Computing Platform of Peking University for enabling our data analysis. We thank the Cloud-Seq Biotech Co. Ltd. (Shanghai) for providing the Ribo-seq service, and we also thank the Emei Tongde Technology Development Co. Ltd (Beijing) for their technical support in single-cell RNA library preparation. This research was funded by the National Natural Science Foundation of China (NSFC 31930016) and the Peking-Tsinghua Center for Life Sciences (to W.W.).

## Author contributions

W.W., X.N., W.T., and Y.L. conceived the project, with W.W. supervising it. W.W., X.N., W.T., Y.L., and Y.-S.L. designed the experiments. X.N. and Y.-S.L. conducted the library screens. X.N. performed the single cell screens, subsequent validations, and experimental data analysis with assistance from B.M.. W.T. handled all bioinformatics analysis. Y.Y. constructed the NGS library. W.T. developed the machine learning model and designed the web page. X.N., W.T., and Y.L. wrote the manuscript, which W.W. revised.

## Competing interests

W.W. is a scientific advisor and founder of EdiGene and Therorna. The remaining authors declare no competing interests.

---

**Extended Data Fig. 1.**
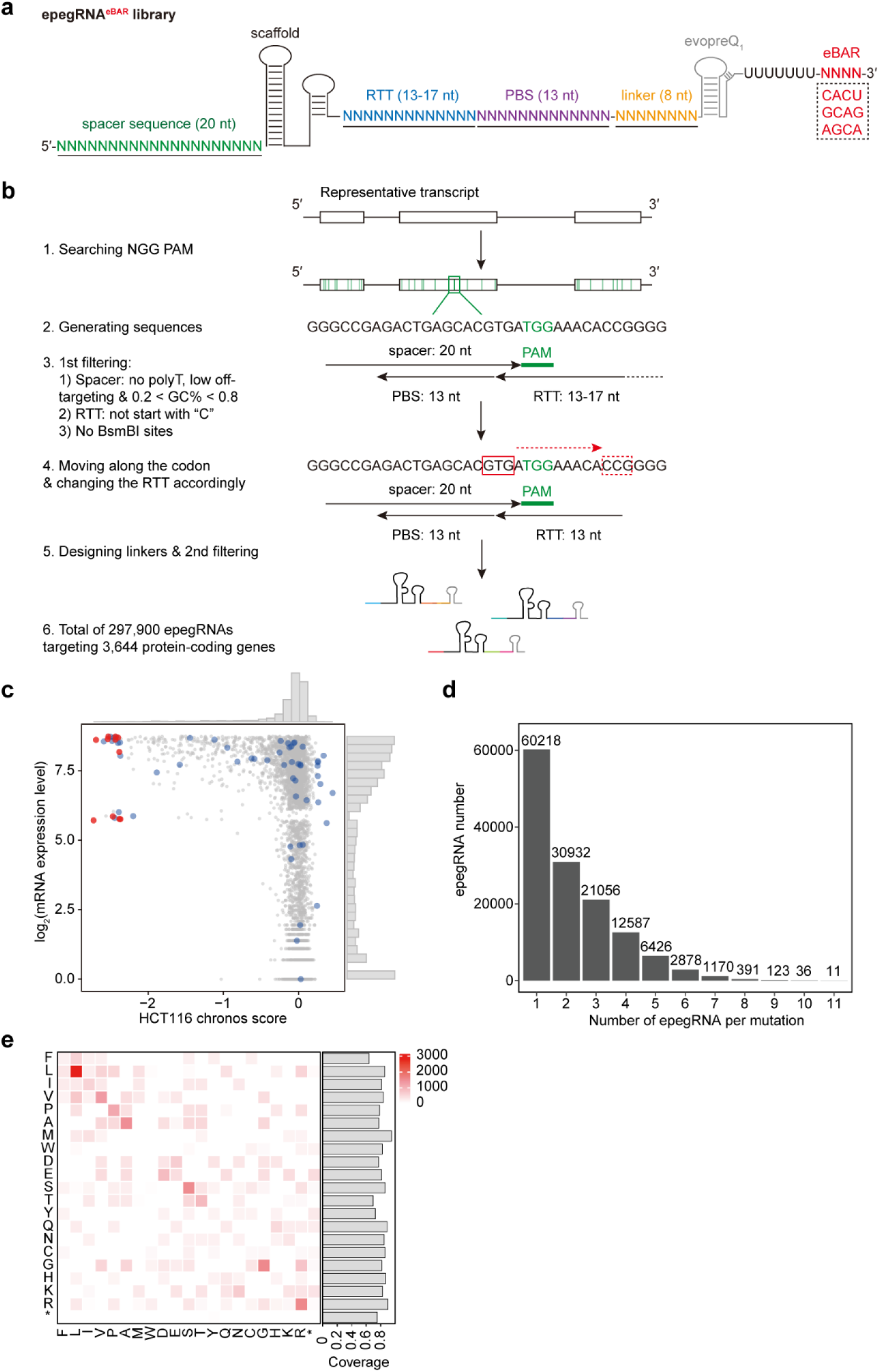
Design of the epegRNA library for screening functional synonymous mutations. **a**, Diagram of the epegRNA^eBAR^ library structure. RTT: reverse transcription template, PBS: primer binding site. **b**, Principles and workflow for designing epegRNAs within the library. Searches for NGG PAMs are conducted on the representative transcript of each gene to determine the spacer sequence, PBS sequence, and RTT, followed by a first filtering step. Comprehensive searches along the coding region are then performed to achieve saturation mutagenesis of synonymous mutations, with linker sequences subsequently designed and a secondary filter applied, resulting in the final epegRNA sequences. **c**, Distribution of expression levels and essentiality of genes within the library in HCT116 cells. Red dots represent the 11 genes targeted for complete saturation mutagenesis, blue dots represent the 56 genes targeted for synonymous saturation mutagenesis, and gray dots represent all other genes. **d**, Relationship between the mutations targeted in the library and the number of corresponding epegRNAs. **e**, Statistics of the types of amino acid substitutions for the 11 genes targeted for saturation mutagenesis. The y-axis represents the original amino acid, the x-axis represents the amino acid post-PE editing, and the values represent the number of epegRNAs designed for each substitution. The histogram on the right illustrates the coverage of the corresponding amino acids, indicating the proportion of amino acids that can be targeted among all amino acids in the 11 genes.

**Extended Data Fig. 2.**
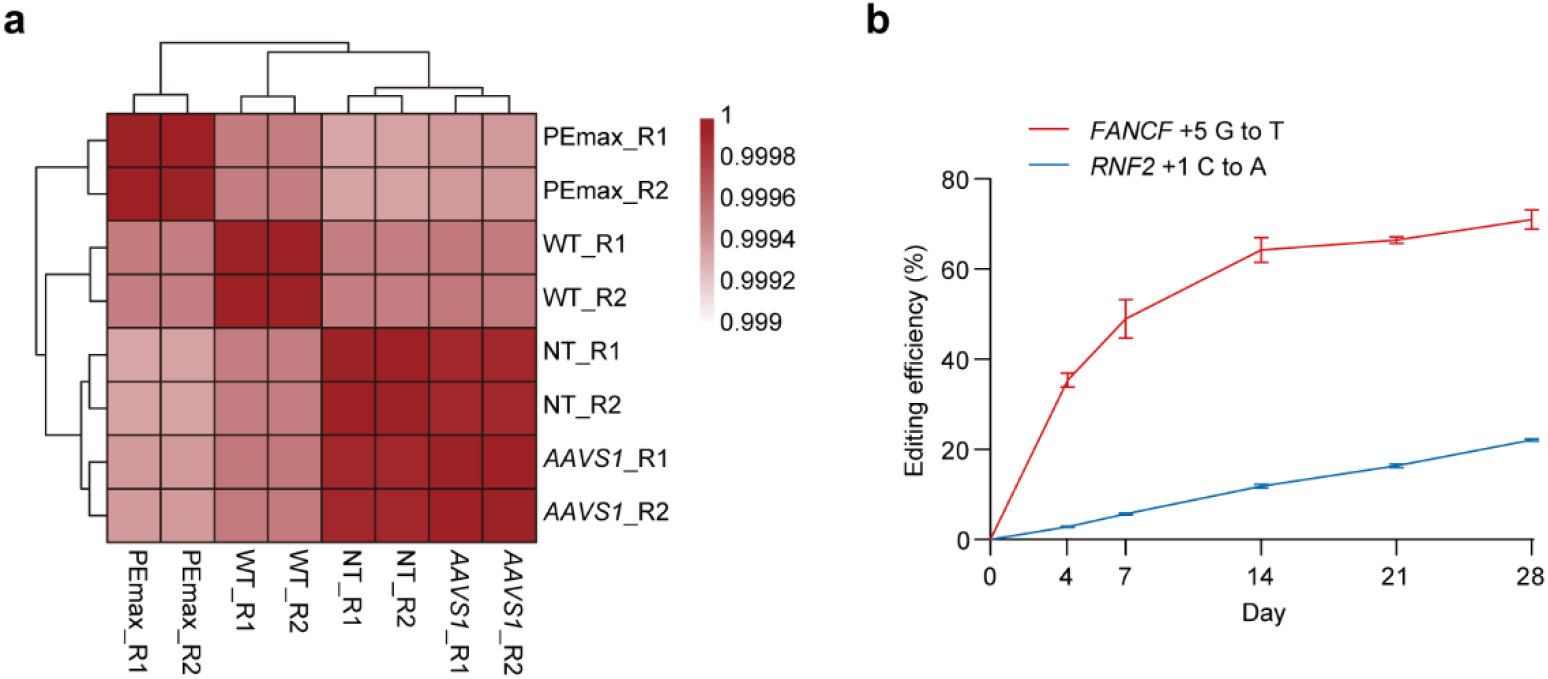
Development of the experimental system using PEmax. **a**, Clustered heatmap displaying Pearson correlation coefficients of the whole transcriptome between HCT116-PEmax cells and wild-type (WT) HCT116 cells, as well as after the addition of nontargeting (NT) epegRNA and *AAVS1*-targeting epegRNA. **b**, Graph showing the changes in editing efficiency over a 28-day period of continuous culture in HCT116-PEmax cells with the addition of two test epegRNAs targeting different sites. The data is presented as the mean ± SD (n = 3).

**Extended Data Fig. 3.**
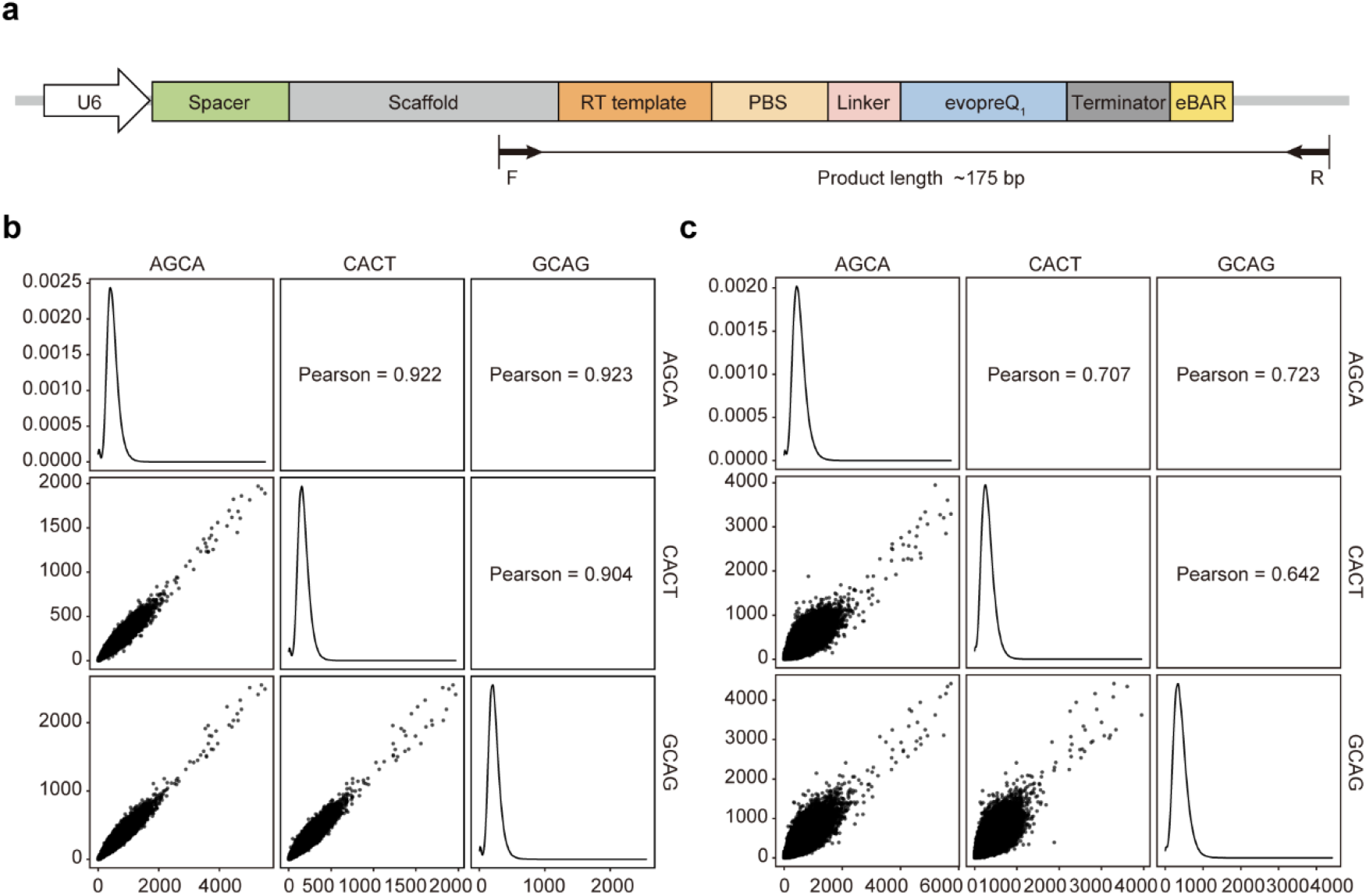
Decoding and data quality control of the epegRNA^eBAR^ library. **a**, Diagram illustrating the genomic PCR process performed on the cell library post-screening. **b**, Correlation among the three internal biological replicates (eBARs) in the library at the start of the experiment (Day 0). **c**, Correlation among the three internal biological replicates (eBARs) in the library at the end of the experiment (Day 35).

**Extended Data Fig. 4.**
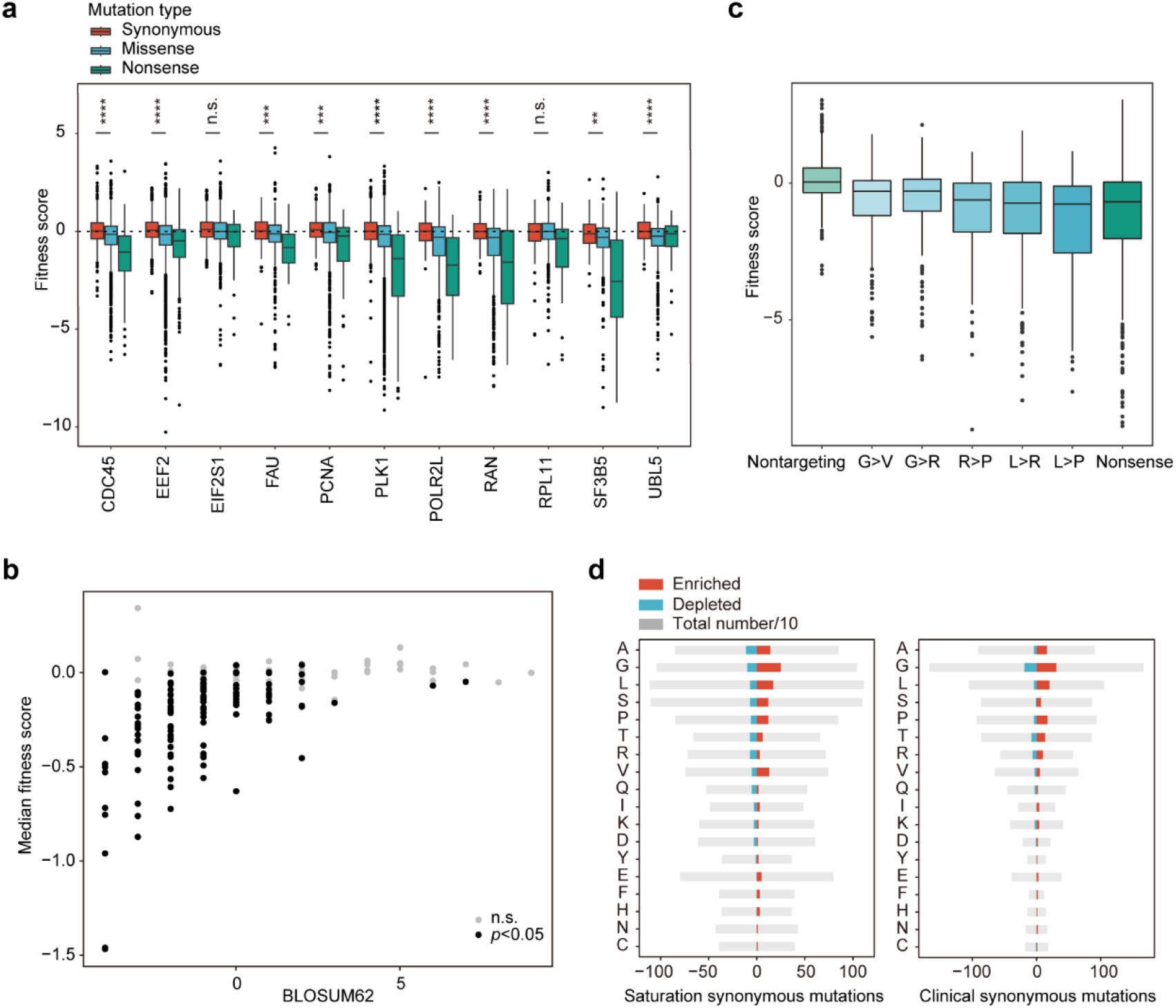
Analysis of characteristics from screen results. **a**, Distribution of fitness scores for the 11 genes subjected to saturation mutagenesis, categorized by mutation type. **b**, Correlation between the BLOSUM62 matrix scores and the screen results for the 11 genes subjected to saturation mutagenesis. In the BLOSUM62 matrix, lower scores indicate lower similarity between two amino acids, reducing likelihood of their substitution. **c**, Identification of several mutation types that significantly impact cell fitness. **d**, Distribution of the number of mutations enriched for various amino acid substitutions in both saturation synonymous mutations and clinical synonymous mutations.

**Extended Data Fig. 5.**
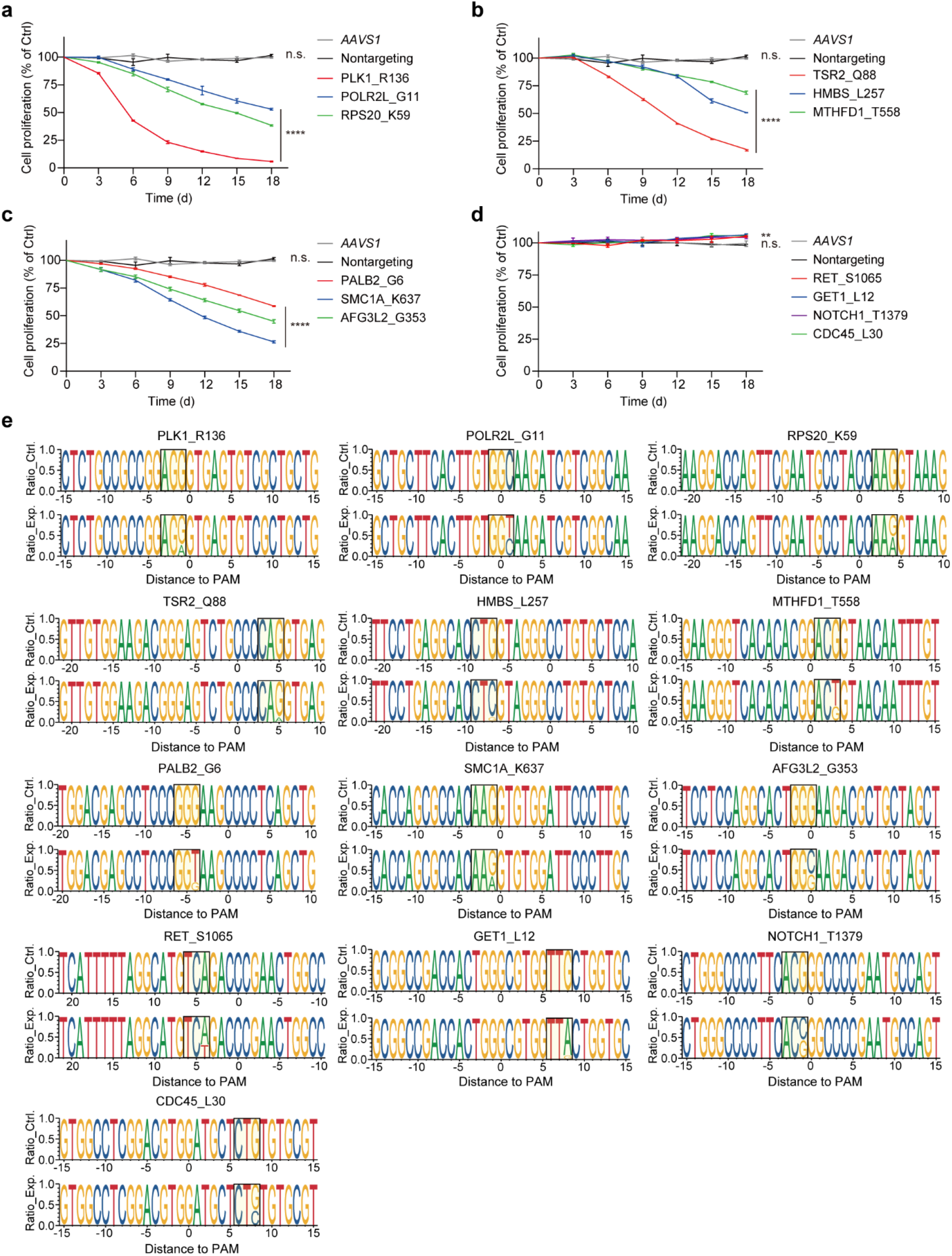
Validation of selected functional synonymous mutations. **a-d**, Cell proliferation rates and **e**, editing results for validated mutation sites. All data are presented as mean ± SD (n = 3). *P* values were calculated using Student’s *t* test, \*\**P* < 0.01, \*\*\*\**P* < 0.0001; n.s., not significant.

**Extended Data Fig. 6.**
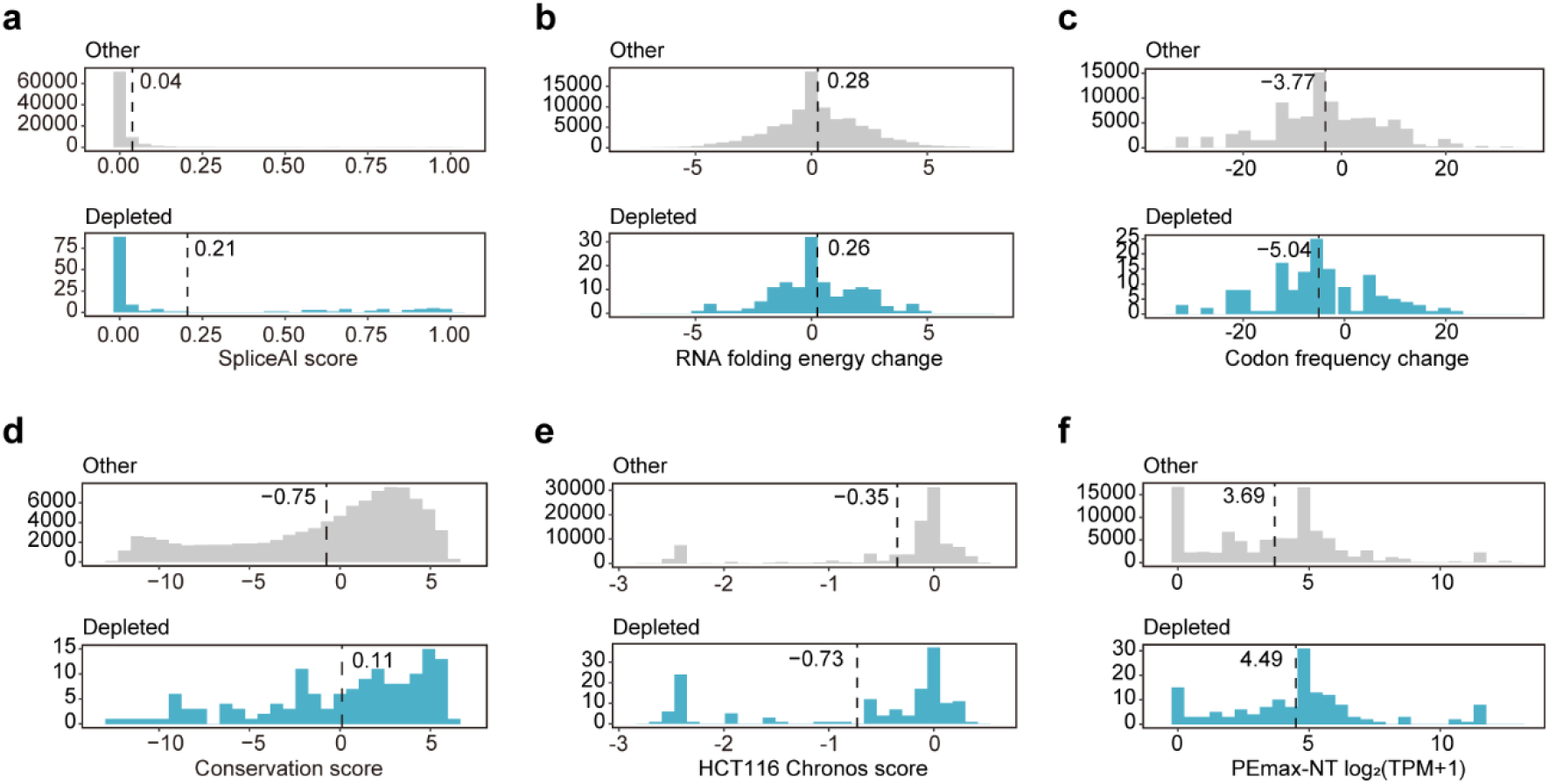
Distribution of key features across synonymous mutation types. **a**, SpliceAI score **b**, Change in RNA folding energy **c**, Change in codon frequency **d**, Conservation score **e**, HCT116 Chronos score and **f**, Gene expression levels.

**Extended Data Fig. 7.**
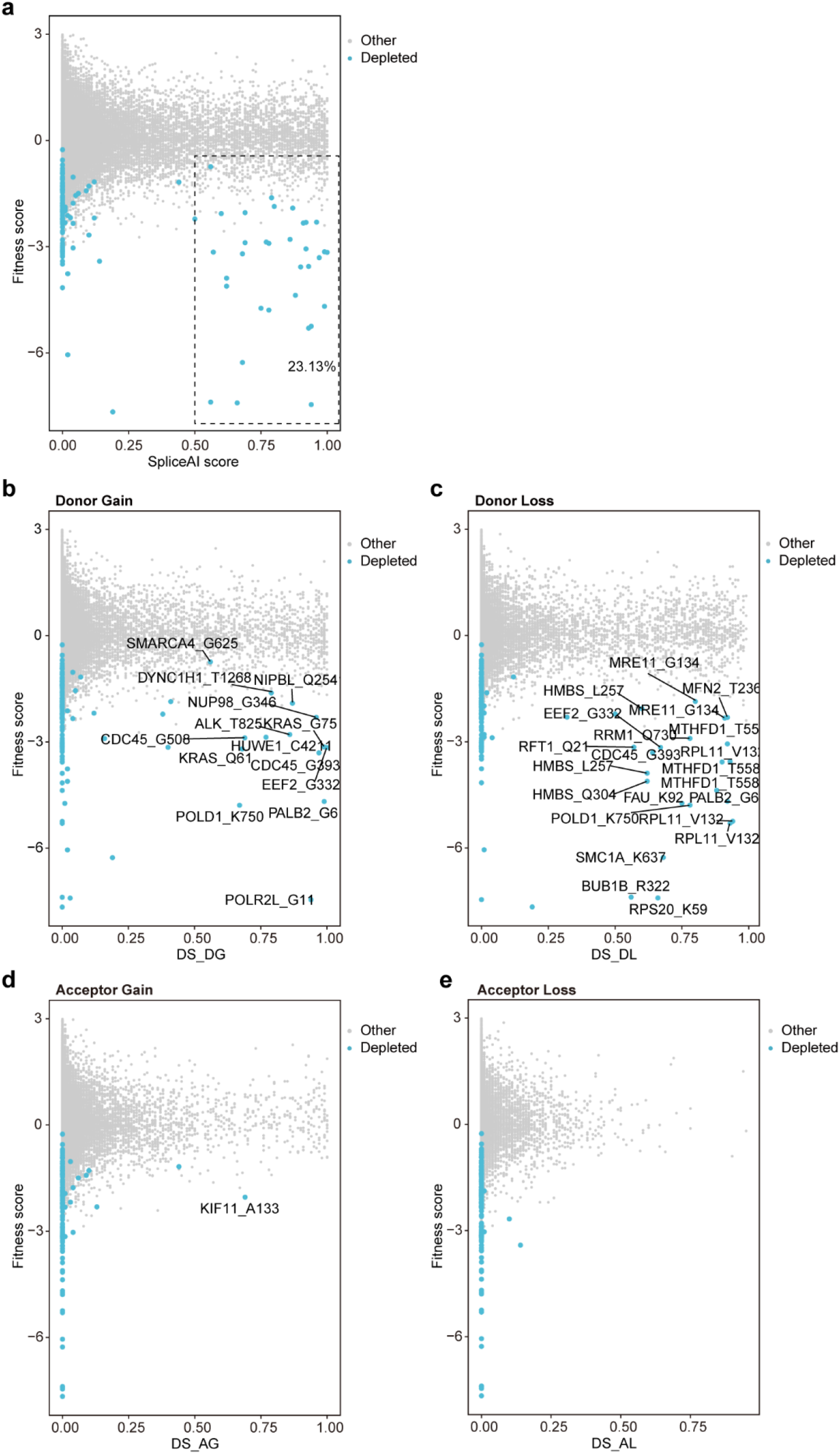
SpliceAI prediction scores and results. **a**, Distribution of splicing scores for mutations in the depletion direction and other non-enriched mutations, using a threshold of 0.5 for higher predictive confidence. **b-e**, SpliceAI scores for synonymous mutations leading to various types of aberrant splicing. DS, Delta score.

**Extended Data Fig. 8.**
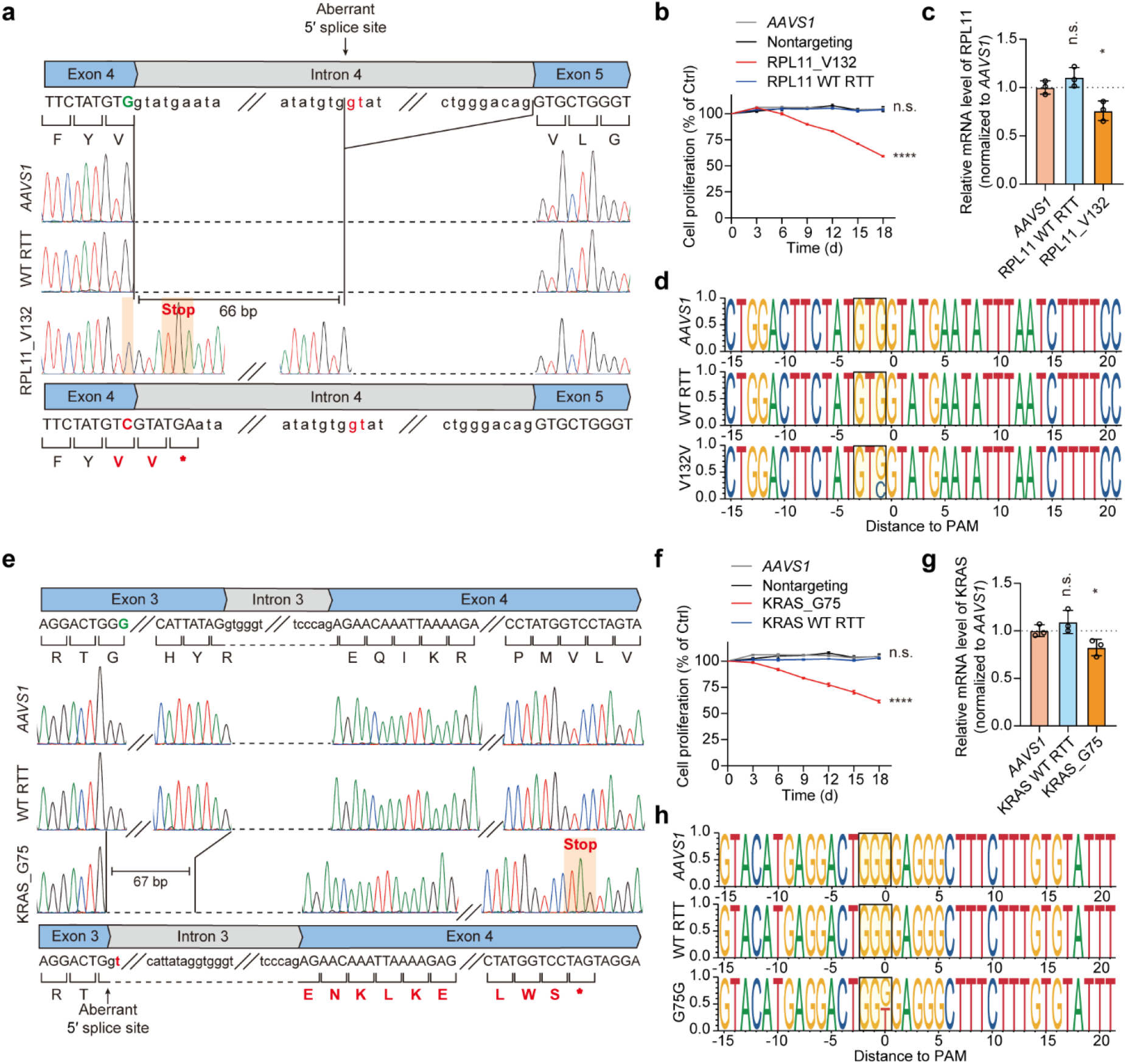
Validation of synonymous mutations causing aberrant RNA splicing. **a**, Schematic depiction of the splicing alterations caused by the RPL11_V132 (GTG>GTC) mutation. **b**, Validation of the effect of the RPL11_V132 (GTG>GTC) mutation on cell proliferation. **c**, Relative mRNA expression levels of *RPL11* in the experimental and control groups. The mRNA level of each sample was quantified by real-time qPCR and normalized by *GAPDH*, and the indicated relative mRNA level was normalized to that of *AAVS1*-targeting control cells. **d**, Analysis of editing outcomes for epegRNA targeting RPL11_V132 and controls via NGS. **e**, Schematic depiction of the splicing alterations caused by the KRAS_G75 (GGG>GGT) mutation. **f**, Validation of the effect of the KRAS_G75 (GGG>GGT) mutation on cell proliferation. **g**, Relative mRNA expression levels of *KRAS* in the experimental and control groups. The mRNA level of each sample was quantified by real-time qPCR and normalized by *GAPDH*, and the indicated relative mRNA level was normalized to that of *AAVS1*-targeting control cells. **h**, Analysis of editing outcomes for epegRNA targeting KRAS_G75 and controls via NGS. All data are presented as mean ± SD (n = 3). *P* values were calculated using Student’s *t* test, \**P* < 0.05, \*\*\*\**P* < 0.0001; n.s., not significant.

**Extended Data Fig. 9.**
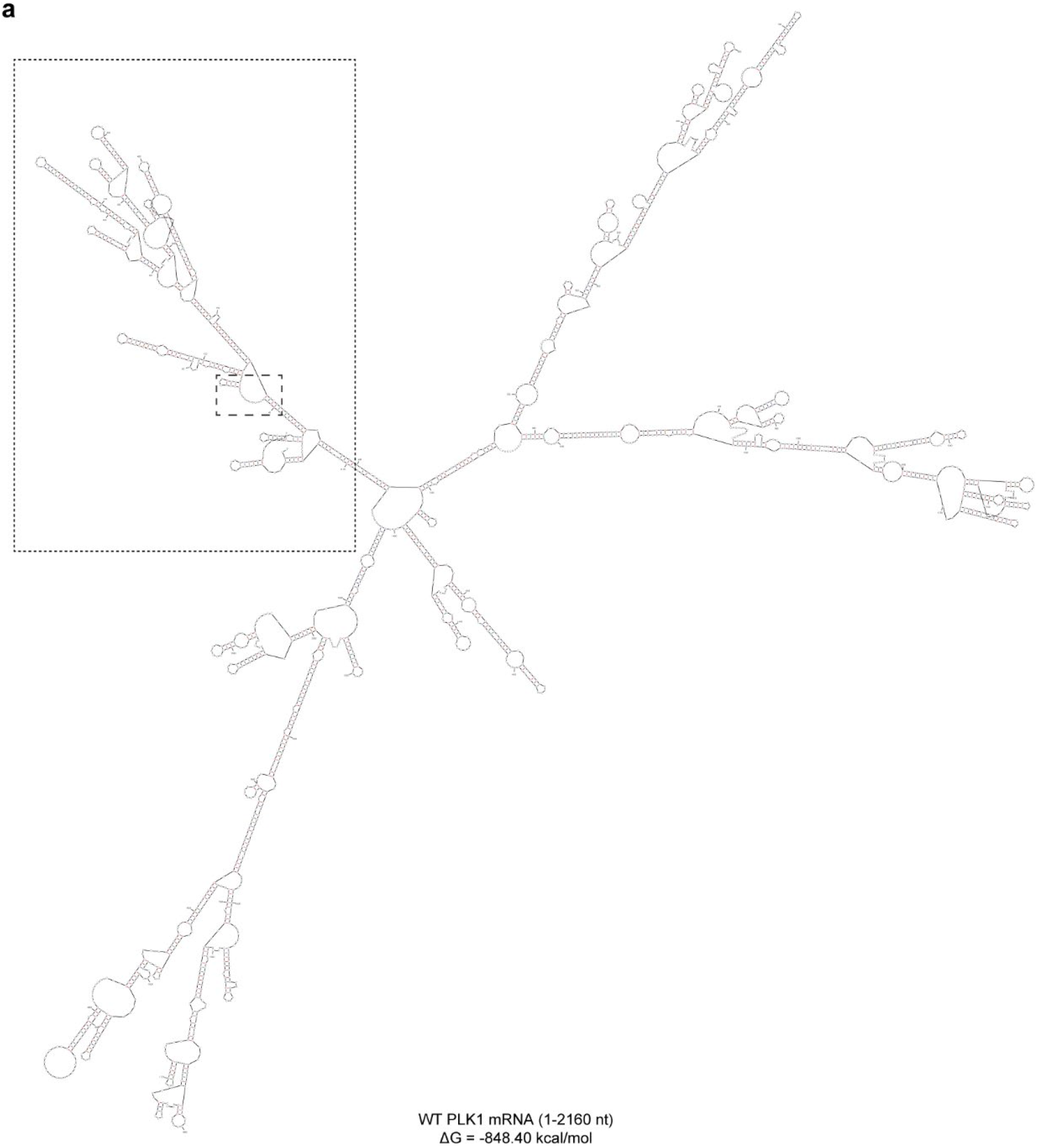
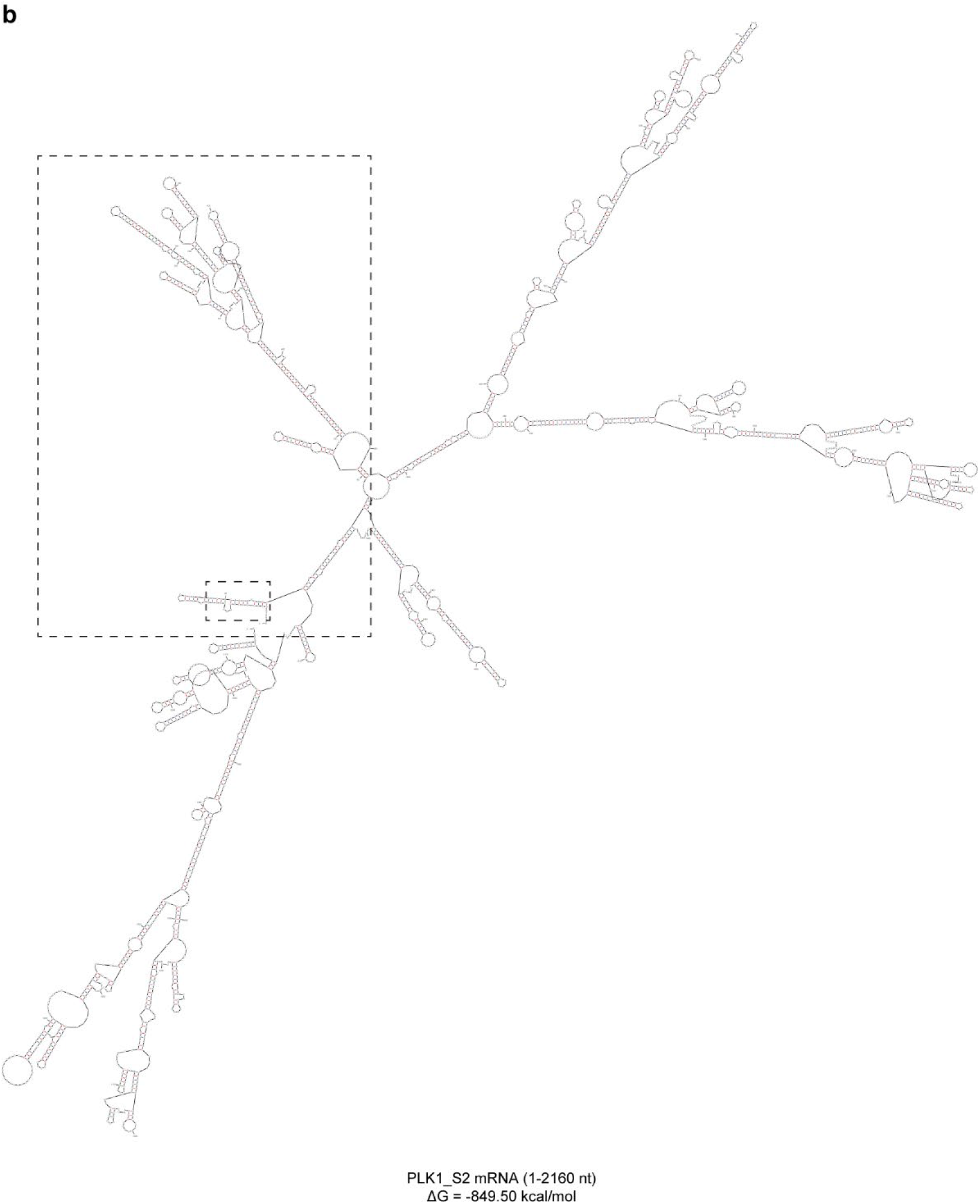
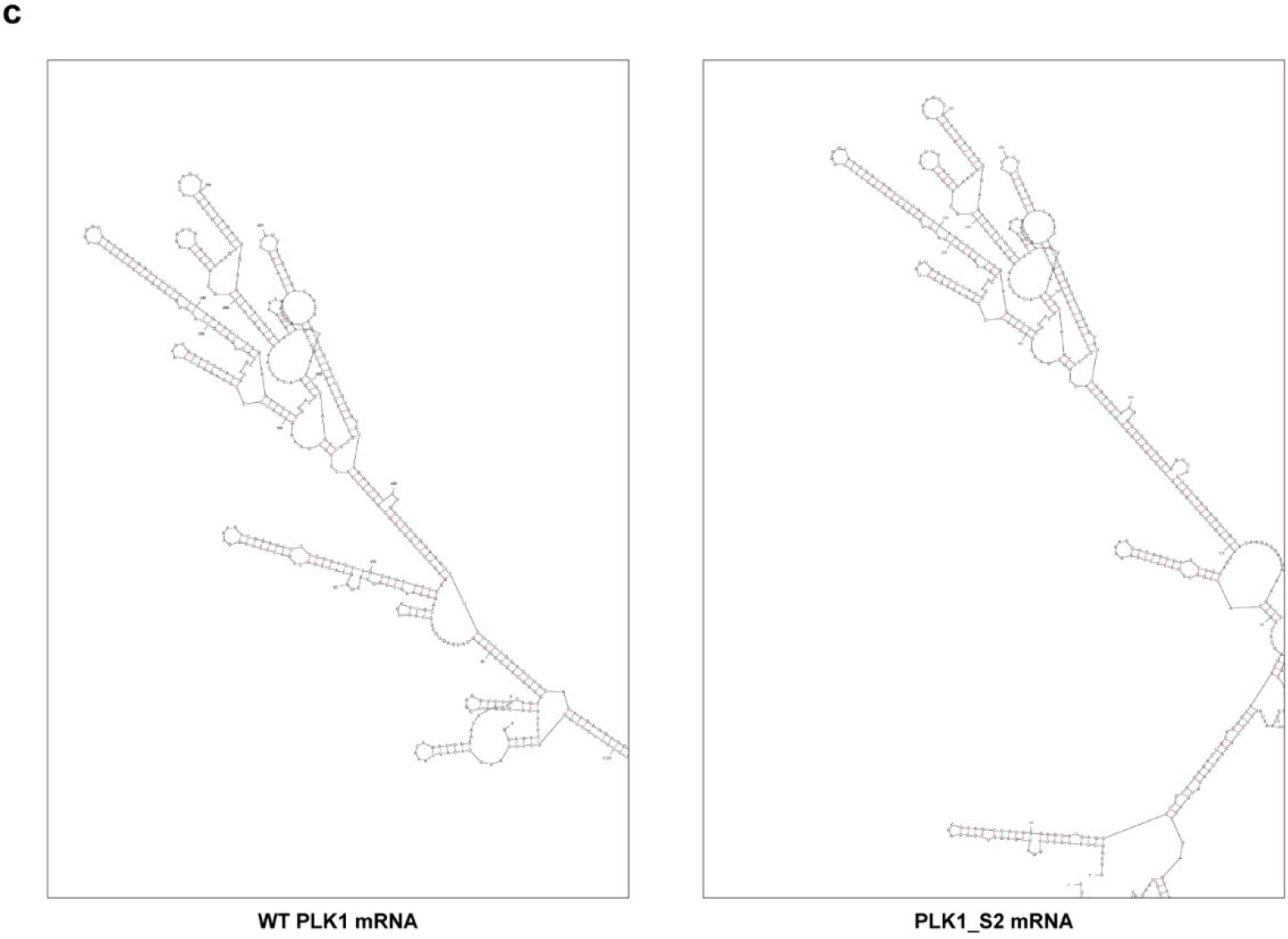
Comparison of *PLK1* mRNA structures before and after the PLK1_S2 (AGT>AGC) mutation. **a**, Structure of wild-type (WT) PLK1 mRNA. **b**, Structure of PLK1_S2 (AGT>AGC) mRNA. **c**, Comparison of the local mRNA structure of *PLK1* before and after the mutation. The secondary structures of mRNA were predicted using RNA Folding Form.

**Extended Data Fig. 10.**
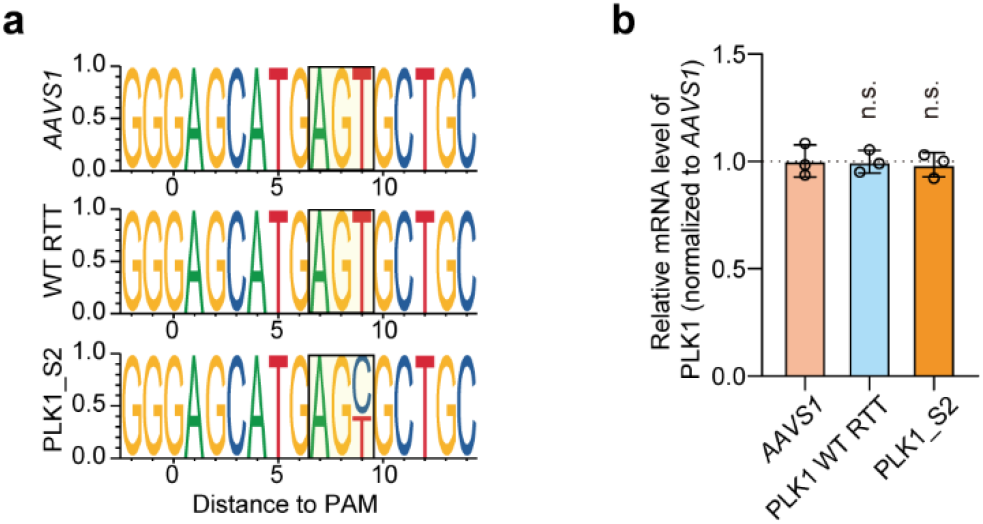
Editing and the gene expression analysis at the PLK1_S2 site. **a**, Analysis of editing outcomes for epegRNA targeting PLK1_S2 and controls via NGS. **b**, Relative mRNA expression levels of *PLK1* in the experimental and control groups. The mRNA level of each sample was quantified by real-time qPCR and normalized by *GAPDH*, and the indicated relative mRNA level was normalized to that of *AAVS1*-targeting control cells. Data are presented as mean ± SD (n = 3). *P* values were calculated using Student’s *t* test, n.s., not significant.

**Extended Data Fig. 11.**
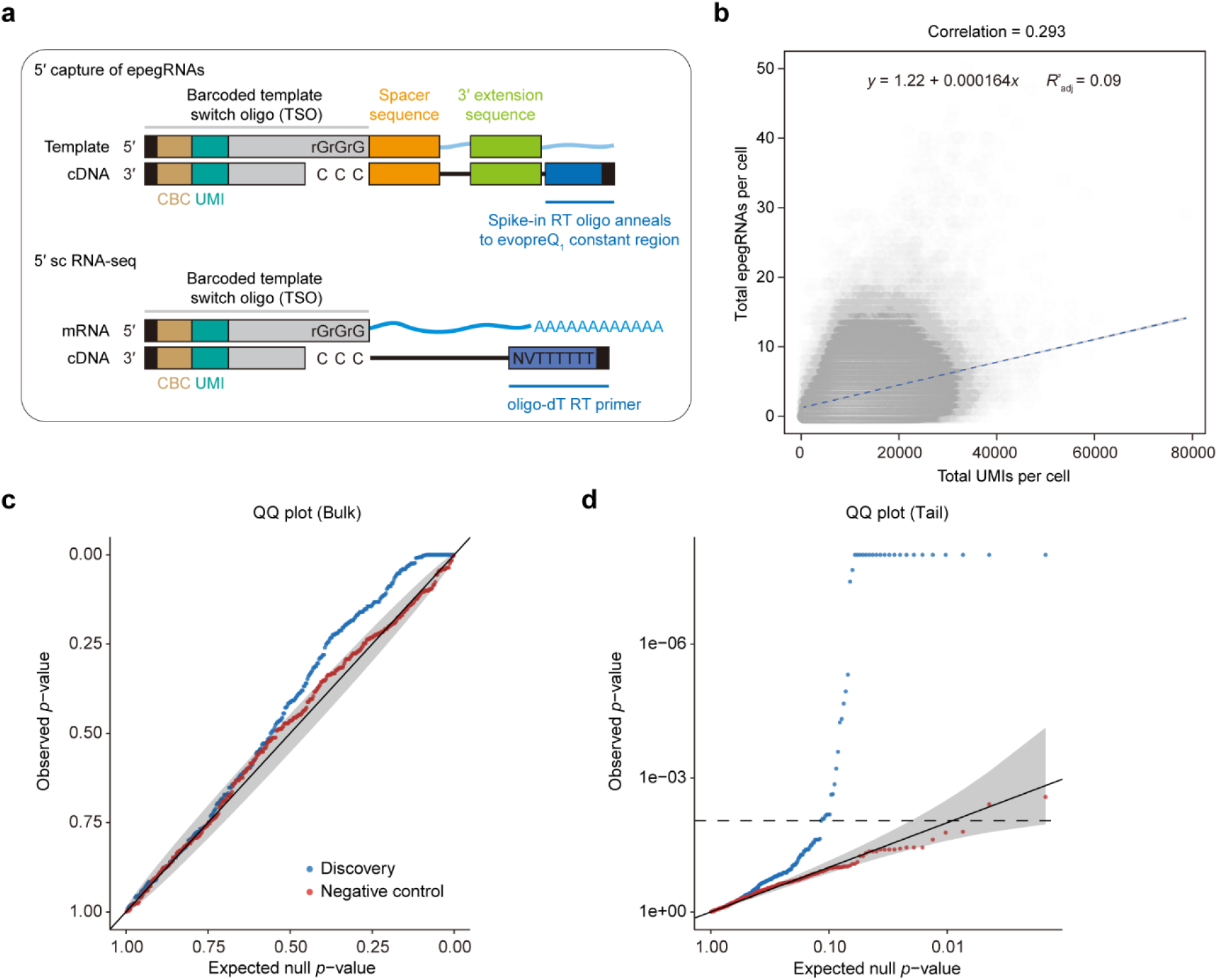
Capture and data quality control for DIRECTED-seq. **a**, Overview of the process for capturing epegRNA and single-cell transcriptomes. **b**, Sequencing depth impact epegRNA detection probability and observed gene expression levels. The blue dashed line represents the linear regression line correlating total epegRNAs per cell (y) with total UMIs per cell (x). **c-d**, Quantile-quantile plots comparing 15 nontargeting epegRNAs (negative control) with 395 epegRNAs across 213 genes (discovery). Genes for the nontargeting tests were randomly selected from the entire gene set.

**Extended Data Fig. 12.**
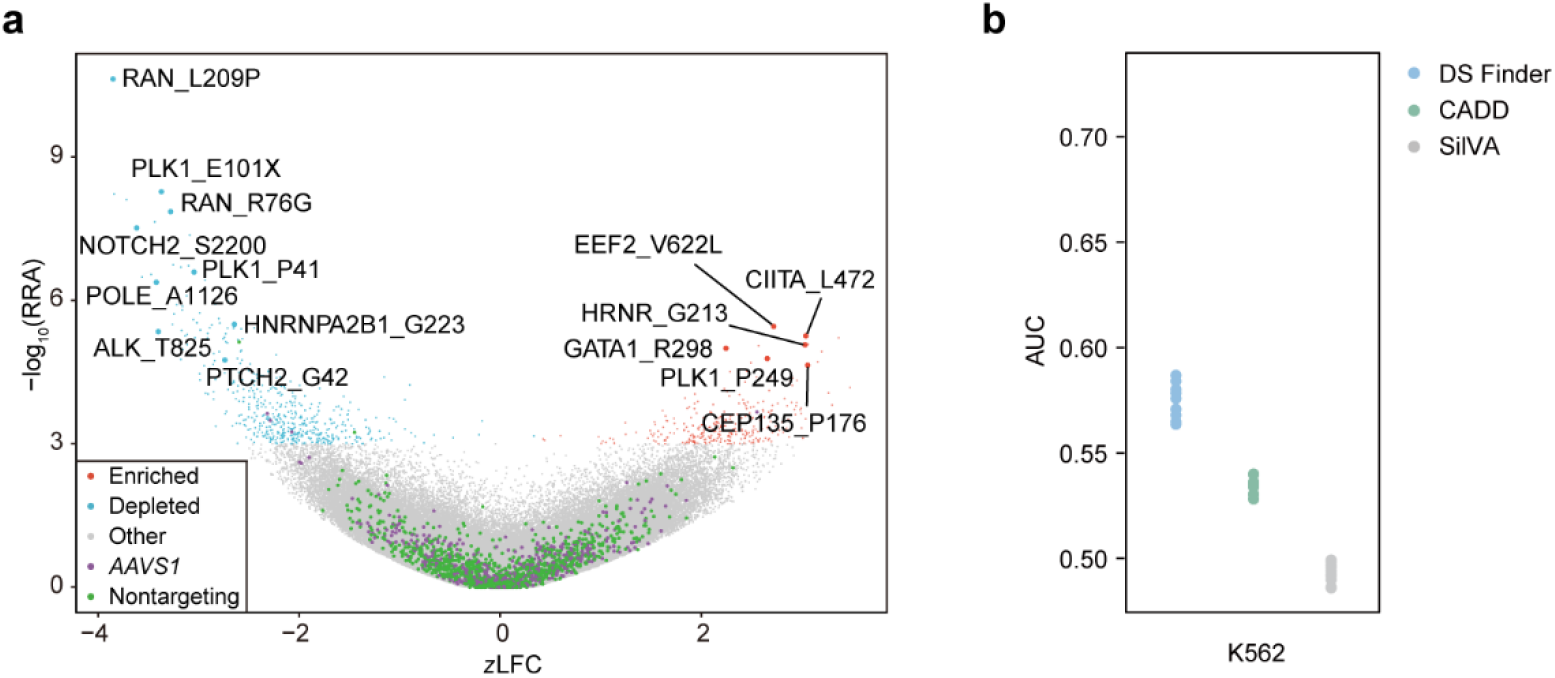
Screening outcomes and model performance in the K562 cell line. **a**, Volcano plot illustrating the results of screening for functional synonymous and nonsynonymous mutations affecting cell fitness. Blue and red dots denote depleted and enriched epegRNAs, respectively. **b**, Performance of DS Finder in the K562 cell line compared with CADD and SilVA. Each point on the graph represents a different dataset, totaling 10.

